# Microbial Communities in Mesopelagic Fish Guts Suggest an Overlooked Component of Marine Biogeochemical Cycles

**DOI:** 10.1101/2025.08.18.670904

**Authors:** Ryan P. Bos, Tracey T. Sutton, Peter R. Girguis

**Author notes:** Corresponding Authors: Ryan P. Bos; Peter R. Girguis. **Competing Interest Statement:** The authors declare no competing interests. **Author Contributions:** R. P. B., T. T. S., and P. R. G. conceptualized the research; R. P. B. and T. T. S. collected samples; R. P. B. and P. R. G. were involved in data generation; R. P. B. and T. T. S. processed the samples; R. P. B. processed the data; R. P. B. analyzed data; R. P. B. wrote the first draft of the manuscript. All authors edited and approved the final version of the manuscript. **Classification:** Biological Sciences >> Environmental Sciences.

## Abstract

Each night, gigatons of oceanic midwater fishes participate in the largest vertical migration on Earth. This movement of organisms from the meso-to the epipelagic and back plays a major role in the flux of organic matter from the surface to the deep sea (the “biological pump”). To date studies have considered the role of fish exports in ocean biogeochemical cycles, but mesopelagic fish gut microbiomes are unquantified and likely play a role in metabolic transformations that influence these cycles. Here, we present data on the abundance of mesopelagic fish gut microbial communities and their functional potential to shape key chemical transformations. Our flow cytometric data from a diversity of taxa reveal that the density of microbes in mesopelagic fish guts is between 10^6^-10^7^ microbes ml^-1^: two to three orders of magnitude higher than the surrounding seawater. In light of the total estimated number of mesopelagic fishes, there are approximately 2.4×10^20^ -1.34×10^24^ gut prokaryotes in total, which without consideration of other biomass abundant animals is realistically within four orders of magnitude of the total prokaryote abundance in mesopelagic waters. Metagenomic analyses revealed distinct bacterial genotypes of the genera *Acinetobacter* and *Psychrobacter* that harbor extensive gene sets for synthesis of essential amino acids, cofactor and vitamins, and short-chain-fatty acids (SCFAs), as well as cellulose and chitin degradation. Given their abundance, mesopelagic fish gut microbiomes are likely playing a role in shaping the nitrogen and carbon cycles, including through calcium carbonate precipitation and sequestration. Moreover, gut microbes are likely more metabolically active than bacterioplankton given that animals actively feed and accumulate organic matter amongst co-concentrated microbes. Gut microbes, especially if we consider all epi-to bathypelagic animals, may represent a significant and previously unrecognized component of marine biogeochemical cycles, though their relative importance compared to free-living microbes requires further quantification.

**Significance Statement:** The daily vertical migration of oceanic midwater fishes represents Earth’s largest animal migration and facilitates the transfer of nutrients from surface to deep waters. Despite their critical role in shaping ocean biogeochemistry, the gut microbiomes of these fishes have remained largely uncharacterized. We demonstrate that mesopelagic fishes harbor dense gut microbial communities -two to three orders of magnitude higher than surrounding seawater-that collectively may be within four orders of magnitude of the total prokaryote abundance in global mesopelagic waters. These gut microbes possess extensive metabolic capabilities for nutrient cycling and likely shape the prevailing biogeochemical and microbial ecological processes in the water column. Given the high likelihood that gut microbes are more metabolically active than free-living bacterioplankton, our findings reveal a previously unrecognized biological component that may fundamentally shape marine biogeochemical cycles and global carbon sequestration.

## Introduction

Deep-pelagic fishes (e.g., those that live in the midwater typically below 200 m depth during daytime) inhabit every ocean basin^1–3^. Many previous studies have worked to estimate total pelagic fish biomass via midwater acoustics, trawl-based surveys, and biomass models^4–13^. These studies have revealed a historic underestimation of midwater biomass in one of the most biodiverse and voluminous habitats on Earth^14,15^. Accounting for 20-31% of the total ocean volume^16–18^, it is likely that the mesopelagic realm (200-1000 m) harbors between 1-33 Gt of fishes alone^4–13^. Realistically, mesopelagic fish biomass is 900% higher than all other fish biomass on Earth^7^ and mesopelagic fish biomass is modelled to increase by 17% by the year 2100 due to warming oceans and shallowing of the deep-scattering layer^17^.

Mesopelagic fishes represent crucial trophic intermediaries as well as important contributors to marine biogeochemical cycles^4^. Every night, colossal assemblages of mesopelagic fishes undertake the largest animal migration on Earth^19^, moving between shallow and deep-pelagic waters to forage under the veil of night. They feed predominantly on chitinous zooplankton containing wax esters^20–24^, organic compounds that sequester 50% of the autotrophic carbon production in the ocean^25^. Upon reaching satiation, these animals return to deeper waters, defecating along the way and expediting the flux of dissolved and particulate organic matter from shallow to deep-pelagic waters^26–29^. Their influence on the flux of organic matter from the surface to the deep sea (the “biological pump”)^30,31^ is pronounced. The extent of active carbon flux by just one family of fishes (the Myctophidae, commonly known as lanternfishes) is thought to be substantial, ranging from 8-28% of passively sinking particulate organic carbon^32–34^. To date, these exports are attributed entirely to the fish and in previous studies there is little consideration of the role of gut microbes in shaping chemical transformations within the digestive tract^35–37^.

It is well known that gut microbes have immense ecophysiological significance in modulating animal health^38–41^, and it is equally likely that they have a similarly massive role in ocean biogeochemical cycles writ large. Gut microbiomes can be thought of as ‘living bioreactors’ that facilitate chemical transformations that either supplement or are beyond the scope of the host’s capabilities, providing access to nutrients that would otherwise be excreted to surrounding seawaters^42–47^. Put a different way, gut microbiomes are co-concentrated microbes and organic matter in small volumes that can expand the breadth of host metabolism and *vice versa*^48,49^. There are most certainly microbially mediated transformations of organic compounds, with varying degrees of recalcitrance within mesopelagic fish guts, that will dictate the nature of organic matter in the fecal mass.

But just how many gut microbes are there in mesopelagic fish guts and what are they capable of doing? To our knowledge, only Ruby and Morin^50^ cultivated and counted 5×10^4^ - 5×10^5^ CFU ml^-1^ bioluminescent bacteria isolated from the deep-sea hatchetfish *Argyropelecus hemigymnus*, but these estimates are certainly a gross underestimation of the total microbial load because the overwhelming majority of microbes will not grow in defined media. Thus, there are no robust estimates of the gut microbial load. Moreover, aside from few amplicon-based sequencing studies and one metagenomic inquiry^51–53^, we still have no genomes from mesopelagic fish gut microbiomes and little insight into their functional metabolic potential. If we use established estimates of median prokaryote abundance (1.51×10^5^ prokaryotes ml^-1^)^54^ in the mesopelagic and posit that the mesopelagic accounts for 20% of ocean volume^16,18^, there are roughly 3.02×10^28^ prokaryotes dispersed in global mesopelagic seawaters^54^. What proportion of microbes living within hosts at mesopelagic depths versus free-living cells, and what their influence is on biogeochemical cycles, remains to be determined.

Here we present flow-cytometric and metagenomic data recovered from a diversity of key mesopelagic fishes, and report on the abundance of fish gut microbial communities and their functional metabolic potential. We used both formalin-fixed and frozen specimens to quantify a range of microbial cells in the guts of ecologically relevant midwater fish species from cosmopolitan deep-pelagic fish families and extrapolate these data to regional and global scales. We focused primarily on determining total microbial cell densities in the guts of diverse vertically migratory and non-migratory taxa with robust documentation via flow cytometry and microscopy to confirm the presence of microbial cells, then provide a first order estimate of microbial abundances in gut microbiomes of highly abundant mesopelagic fishes. As there are limited metagenomic data for mesopelagic fish gut microbiomes, we also performed whole-genome-shotgun sequencing followed by a granular annotation scheme to explore potential activities of this heretofore recognized microbial reservoir.

## Results

### Flow Cytometry

Formalin-fixed samples from the Harvard Museum of Comparative Zoology (MCZ) were classified as “museum” samples, whereas frozen samples collected from the Gulf of Mexico were classified as “frozen” samples. A total of 98 museum (13-58 years) and 24 frozen (2-3 years) mesopelagic fishes from four cosmopolitan families were processed on the Aurora™ flow cytometer (Figure 1; Tables 1 and S1). Representative flow cytometric scatter plots and microbe images are displayed in Figure 2. Staining of gut supernatants followed by flow cytometry and hierarchical gating of only cells less than three microns revealed a mean of 2.01×10^7^ ± 2.15×10^6^ SE microbes ml^-1^ across museum fish samples, with a range of 5.8×10^5^ – 7.8×10^7^ microbes ml^-1^. Standard lengths ± SD, fish wet masses ± SD, wet mass of guts ± SD, and microbial load ± SD for each fish species can be viewed in Table 1. Standard length (Spearman’s correlation, ρ = 0.45), fish wet mass (Spearman’s correlation, ρ = 0.56), and wet mass gut (Spearman’s correlation, ρ = 0.49) were moderately positively correlated with microbes ml^-1^.

**Figure 1.**
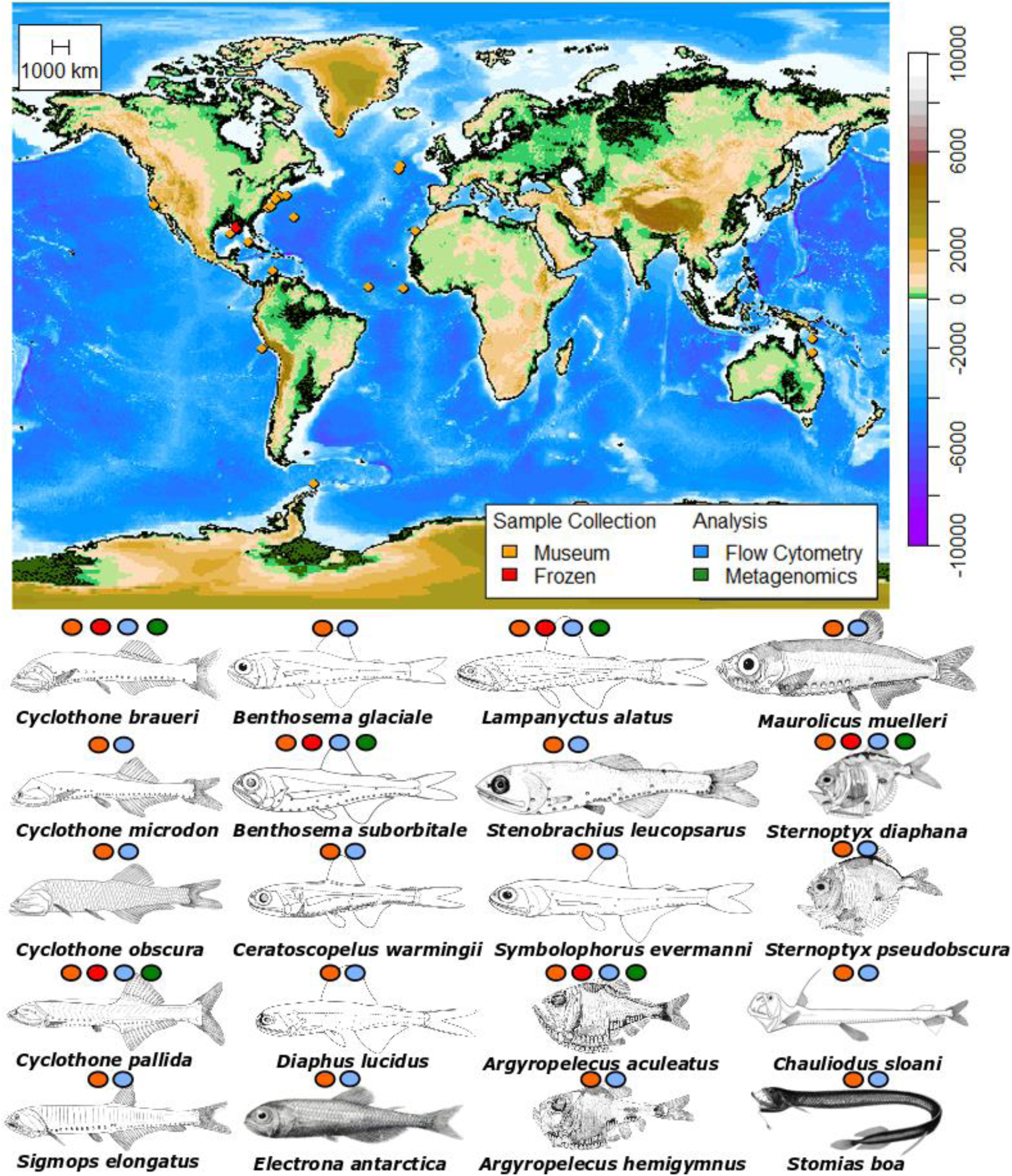
Mercator projection of original locations of frozen (red) and museum (orange) mesopelagic fish collections and species used in either flow cytometric (blue) and metagenomic (green) analyses. See Table S1 for more details.

**Figure 2.**
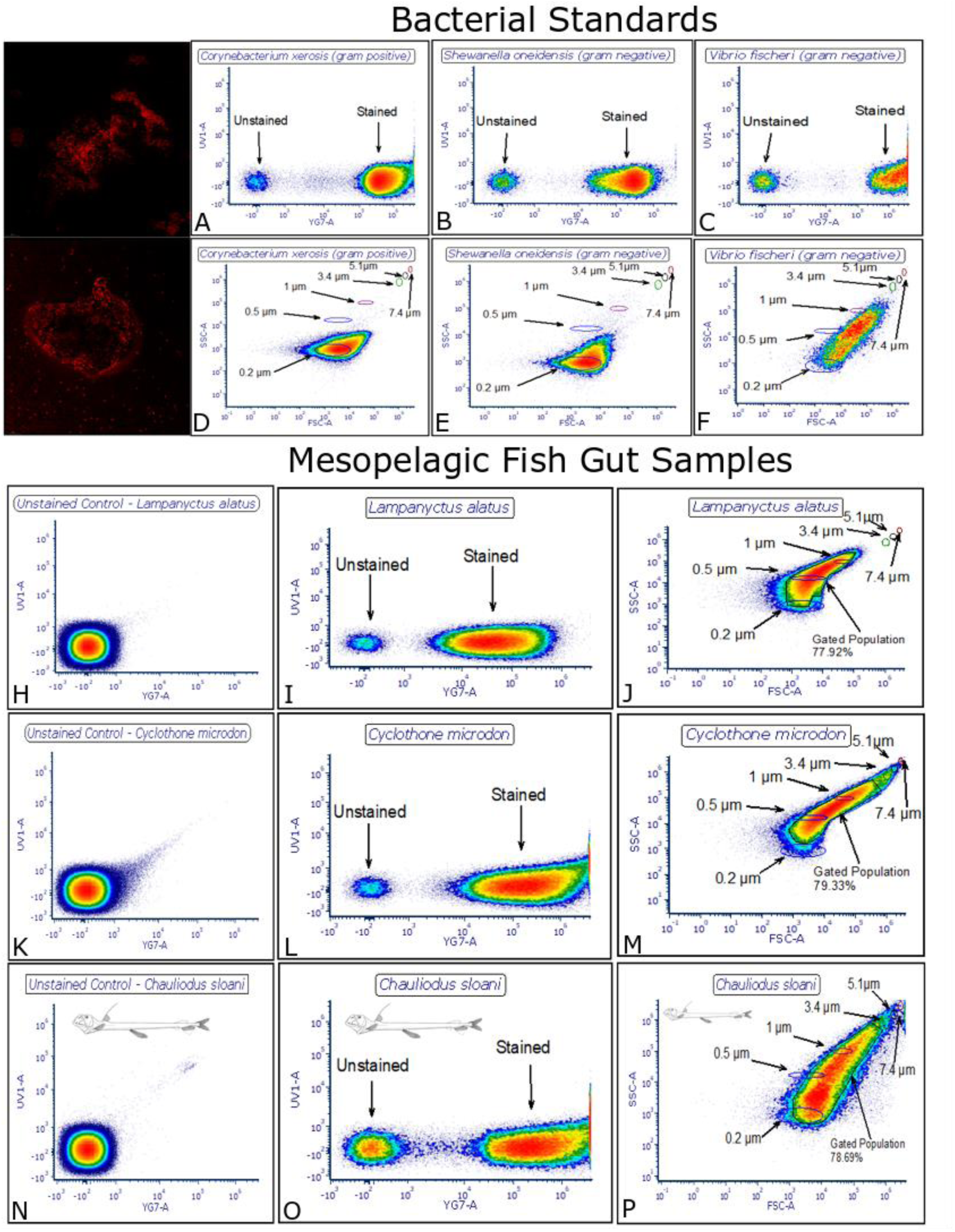
Typical stained microbial cells from mesopelagic fish guts (top left panel). Representative scatter plots of unstained controls (H, K, N), fluorescent intensity of unstained and stained cellular populations (A-C, I, L, O), and hierarchically gated FSC-A and SSC-A (D-F, J, M, P) for bacterial standards (top panel) and fish gut microbiomes (bottom panel) using Aurora flow cytometry.

**Table 1.** The fish wet mass (g) ± SD, fish gut wet mass (g) ± SD, microbial cells ml^-1^ ± SD, migration behavior, and feeding guild of mesopelagic fish species from museum samples of cosmopolitan deep-pelagic families that were utilized for flow cytometric analyses. VM = vertical NVM = non-vertical migrator. Refer to Bos et al.^107^ for more information regarding depths used for migration classifications.

| Species | Standard Length (mm) ± SD | Fish Wet Mass (g) ± SD | Fish Gut Wet Mass (g) ± SD | Microbial Cells ml<sup>-1</sup> ± SD (Aurora) | Migration Behavior | Feeding Guild | Sample Size (n) |
| --- | --- | --- | --- | --- | --- | --- | --- |
| **Gonostomatidae** |  |  |  |  |  |  |  |
| *Cyclothone braueri* | 18.59 - 23.45 ± 2.14 | 0.0407 - 0.0801 ± 0.02 | 0.0016 - 0.0493 ± 0.01862 | 5.8x10<sup>5</sup> - 1.98x10<sup>6</sup> ± 4.99x10<sup>5</sup> | NVM | Detritivore, Gelatinovore, Zooplanktivore | 5 |
| *Cyclothone microdon* | 41.42 - 50.62 ± 3.89 | 0.3286 - 0.8317 ± 0.19 | 0.021 - 0.0611 ± 0.01627 | 6.96x10<sup>5</sup> - 4.94x10<sup>6</sup> ± 1.46x10<sup>6</sup> | NVM | Detritivore, Gelatinovore, Zooplanktivore | 5 |
| *Cyclothone obscura* | 39.4 - 49.2 ± 3.97 | 0.2851 - 0.5408 ± 0.09 | 0.0191 - 0.0814 ± 0.03 | 2.23x10<sup>6</sup> - 1.1x10<sup>7</sup> ± 3.32x10<sup>6</sup> | NVM | Detritivore, Gelatinovore, Zooplanktivore | 5 |
| *Cyclothone pallida* | 32.71 - 45.36 ± 5.50 | 0.1695 - 0.4575 ± 0.13 | 0.00741 - 0.0291 ± 0.01 | 4.7x10<sup>6</sup> - 2.69x10<sup>7</sup> ± 7.44x10<sup>6</sup> | NVM | Detritivore, Gelatinovore, Zooplanktivore | 5 |
| *Sigmops elongatus* | 71.05 - 167.6 ± 37.28 | 1.15 - 14.42 ± 5.76 | 0.0453 - 0.5029 ± 0.22 | 2.53x10<sup>7</sup> - 6.4x10<sup>7</sup> ± 1.38x10<sup>7</sup> | VM | Piscivore, Zooplanktivore | 5 |
| **Myctophidae** |  |  |  |  |  |  |  |
| *Benthoosema glaciale* | 34.59 - 56.19 ± 7.77 | 0.6421 - 2.8146 ± 0.82 | 0.0145 - 0.1017 ± 0.04 | 2.93x10<sup>6</sup> - 1.08x10<sup>7</sup> ± 2.84x10<sup>6</sup> | VM | Zooplanktivore | 5 |
| *Benthoosema suborbitale* | 20.72 - 24.14 ± 1.45 | 0.1224 - 0.1905 ± 0.03 | 0.0039 - 0.1489 ± 0.06 | 5.81x10<sup>5</sup> - 4.75x10<sup>7</sup> ± 1.74x10<sup>7</sup> | VM | Zooplanktivore | 5 |
| *Ceratoscopelus warmingii* | 32.53 - 56.53 ± 9.78 | 0.4975 - 2.2965 ± 0.75 | 0.0037 - 0.0635 ± 0.03 | 1.33x10<sup>6</sup> - 3.66x10<sup>7</sup> ± 1.43x10<sup>7</sup> | VM | Zooplanktivore | 5 |
| *Diaphus lucidus* | 20.91 - 55.64 ± 15.12 | 0.193 - 2.8041 ± 1.19 | 0.0003 - 0.0887 ± 0.04 | 1.47x10<sup>6</sup> - 4.11x10<sup>7</sup> ± 1.44x10<sup>7</sup> | VM | Zooplanktivore | 5 |
| *Electrona antarctica* | 49.82 - 78.63<br>± 11.58 | 1.7069 - 10.4503 ± 3.54 | 0.0711 - 0.3506 ± 0.13 | 2.87x10<sup>7</sup> -<br>7.68x10<sup>7</sup> ±<br>2.19x10<sup>7</sup> | VM | Zooplanktivore | 5 |
| *Lampanyctus alatus* | 37.03 - 44.22<br>± 2.89 | 0.5761 - 0.8881 ± 0.14 | 0.0148 - 0.1107 ± 0.04 | 7.22x10<sup>5</sup> -<br>3.72x10<sup>6</sup> ±<br>1.14x10<sup>6</sup> | VM | Zooplanktivore | 5 |
| *Stenobrachius leucopsarus* | 43.02 - 68.22<br>± 9.59 | 0.9926 - 3.8724 ± 1.17 | 0.0872 - 0.353 ± 0.13 | 1.27x10<sup>6</sup> -<br>3.3x10<sup>7</sup> ±<br>1.21x10<sup>7</sup> | VM | Zooplanktivore | 5 |
| *Symbolophorus evermanni* | 66.66 - 83.49<br>± 7.74 | 4.5925 - 7.985 ± 1.57 | 0.1201 - 0.2424 ± 0.05 | 1.05x10<sup>7</sup> -<br>4.26x10<sup>7</sup> ±<br>1.19x10<sup>7</sup> | VM | Zooplanktivore | 4 |
| **Sternoptychidae** |  |  |  |  |  |  |  |
| *Argyropelecus aculeatus* | 45.31 - 67.04<br>± 9.06 | 3.5685 - 13.453 ± 4.16 | 0.1166 - 1.6179 ± 0.71 | 3.59x10<sup>7</sup> -<br>7.02x10<sup>7</sup> ±<br>1.55x 10<sup>7</sup> | VM | Generalist | 4 |
| *Argyropelecus hemigymnus* | 23.19 - 29.22<br>± 2.51 | 0.5277 - 1.211 ± 0.28 | 0.008 - 0.048 ± 0.02 | 1.6x10<sup>6</sup> -<br>2.52x10<sup>7</sup> ±<br>8.27x10<sup>6</sup> | VM | Generalist | 5 |
| *Maurollicus muelleri* | 41.7 - 48.73<br>± 2.90 | 0.8305 - 1.772 ± 0.37 | 0.0112 - 0.4055 ± 0.17 | 2.3x10<sup>7</sup> -<br>7.8x10<sup>7</sup> ±<br>1.78x10<sup>7</sup> | VM | Zooplanktivore | 5 |
| *Sternoptyx diaphana* | 15.7 - 29.51<br>± 5.36 | 0.3267 - 1.8363 ± 0.59 | 0.0527 - 0.0814 ± 0.01 | 8.18x10<sup>6</sup> -<br>2.89x10<sup>7</sup> ±<br>7.45x10<sup>6</sup> | NVM | Generalist | 5 |
| *Sternoptyx pseudobscura* | 19.35 - 26.08<br>± 2.65 | 0.7105 - 2.1197 ± 0.57 | 0.1296 - 0.4989 ± 0.16 | 2.19x10<sup>6</sup> -<br>5.2x10<sup>7</sup> ±<br>1.83x10<sup>7</sup> | NVM | Generalist | 5 |
| **Stomiidae** |  |  |  |  |  |  |  |
| *Chauliodus sloani* | 32.53 -<br>192.69 ±<br>58.26 | 1.734 - 14.98 ± 6.14 | 0.1181 - 1.0626 ± 0.44 | 2.15x10<sup>6</sup> -<br>2.33x10<sup>7</sup> ±<br>8.77x10<sup>6</sup> | VM | Piscivore | 4 |
| *Stomias boa* | 111.77 - 239<br>± 51.84 | 4.9833 - 32.614 ± 10.37 | 0.2199 - 1.4235 ± 0.53 | 6.66x10<sup>6</sup> -<br>5.57x10<sup>7</sup> ±<br>1.56x10<sup>7</sup> | VM | Piscivore | 5 |

When separating museum fishes at the family taxonomic level, Gonostomatidae, Myctophidae, Sternoptychidae, and Stomiidae harbored a mean of 1.01×10^7^ ± 1.13×10^7^, 1.73×10^7^ ± 2.25×10^7^, 2.7×10^7^ ± 2.11×10^7^, and 4.15×10^7^ ± 1.38×10^7^ microbes ml^-1^, respectively. When comparing microbes ml^-1^ at the family level for museum fishes, there was a statistical difference between families (Kruskal-Wallis, *p* = 0.00018). Sternoptychidae and Stomiidae had significantly higher microbes ml^-1^ than Gonostomatidae, and Stomiidae also had significantly more cells when compared with Myctophidae (Figure 3A). When binning museum fishes by vertical migration behavior (i.e., migratory, non-migratory), there was no statistical difference in microbes ml^-1^ (Mann-Whitney, *p* = 0.1315, *W =* 1161, Figure 3B). When comparing normalized gut microbial counts between frozen and museum fish species (Figure 3C), five of the six species compared, *Argyropelecus aculeatus* (Mann-Whitney, *W* = 10, *p* = 0.6857), *Argyropelecus hemigymnus* (Mann-Whitney, *W* = 16, *p* = 0.1905), *Benthosema suborbitale* (Mann-Whitney, *W* = 10, *p* = 1), *Cyclothone obscura* (Mann-Whitney, *W* = 8, *p* = 0.7302), and *Lampanyctus alatus* (Mann-Whitney, *W* = 2, *p* = 0.06349) harbored statistically similar microbes ml^-1^ g^-1^, whereas *Cyclothone pallida* (Mann-Whitney, *W* = 20, *p* = 0.01587) had significantly higher microbes ml^-1^ g^-1^ in museum samples.

**Figure 3.**
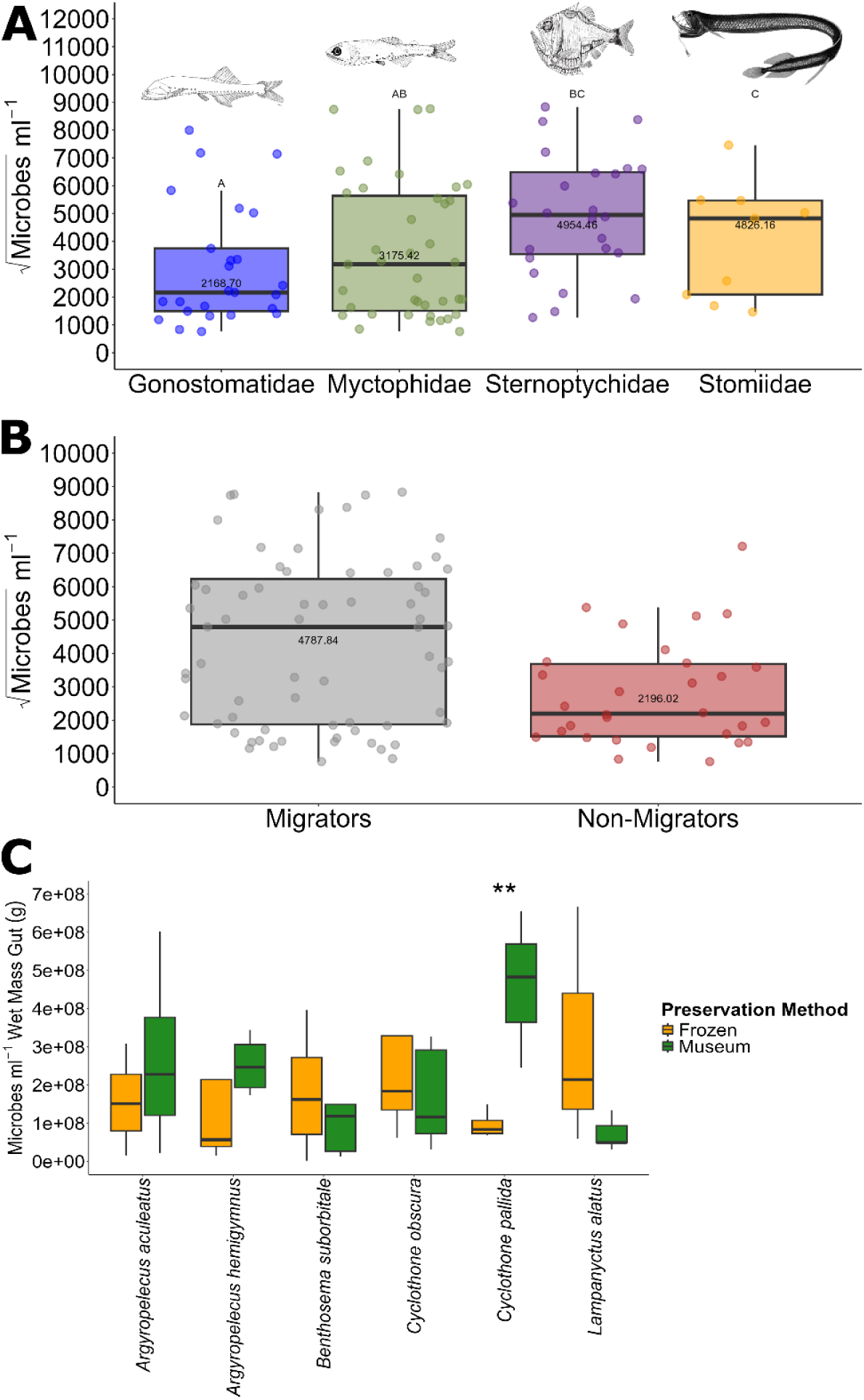
A) Boxplots containing microbes ml^-1^ (square root transformed for resolution) from museum deep-pelagic gut microbiomes separated at the family taxonomic level. B) Boxplots containing microbes ml^-1^ (square root transformed for resolution) from museum deep-pelagic fish guts separated into migrators and non-migrators. Boxes represent the interquartile range and solid horizontal black line is the median. Letters (A, B, C) represent statistical groupings from post hoc multiple comparisons. Numbers directly above the median are the median value for each group. Square root transformation is for graphical clarity. C) Boxplots comparing standardized microbes ml^-1^ between select species that were frozen or archived museum samples. ** denotes statistical significance between microbes ml^-1^.

Global mesopelagic fish biomass estimates^4^ (g m^-2^) from defined Food and Agricultural Organization (FAO) regions are displayed in Figure S1. Using the above biomass data (1 Gt), the 11 different scenarios of the average mass of a mesopelagic fish (0.5 g, 1 g, 2 g, 3 g, 4 g, 5 g, 6 g, 7 g, 8 g, 9 g, 10 g; essentially one order of magnitude), and the geometric mean of gut microbes from museum specimens observed in this study (2.01×10^7^ microbes ml^-1^), the resulting estimate of regional and global mesopelagic fish and fish gut microbes was calculated and is displayed in Figure 4. For a conservative 1 Gt of biomass, the global mesopelagic fish count ranged from 1.67×10^14^ to 2×10^15^ fishes, whereas global fish gut microbe count spanned from 2.4×10^20^ – 2.88×10^21^. In addition, recently updated acoustic mesopelagic fish biomass estimates suggest there are up to 33 Gt (also presented in Table 2). When considering the recent biomass estimates, there is potentially a range of 1.0×10^14^ – 6.66×10^16^ fishes harboring 2.4×10^20^ -1.34×10^24^ gut prokaryotes.

**Figure 4.**
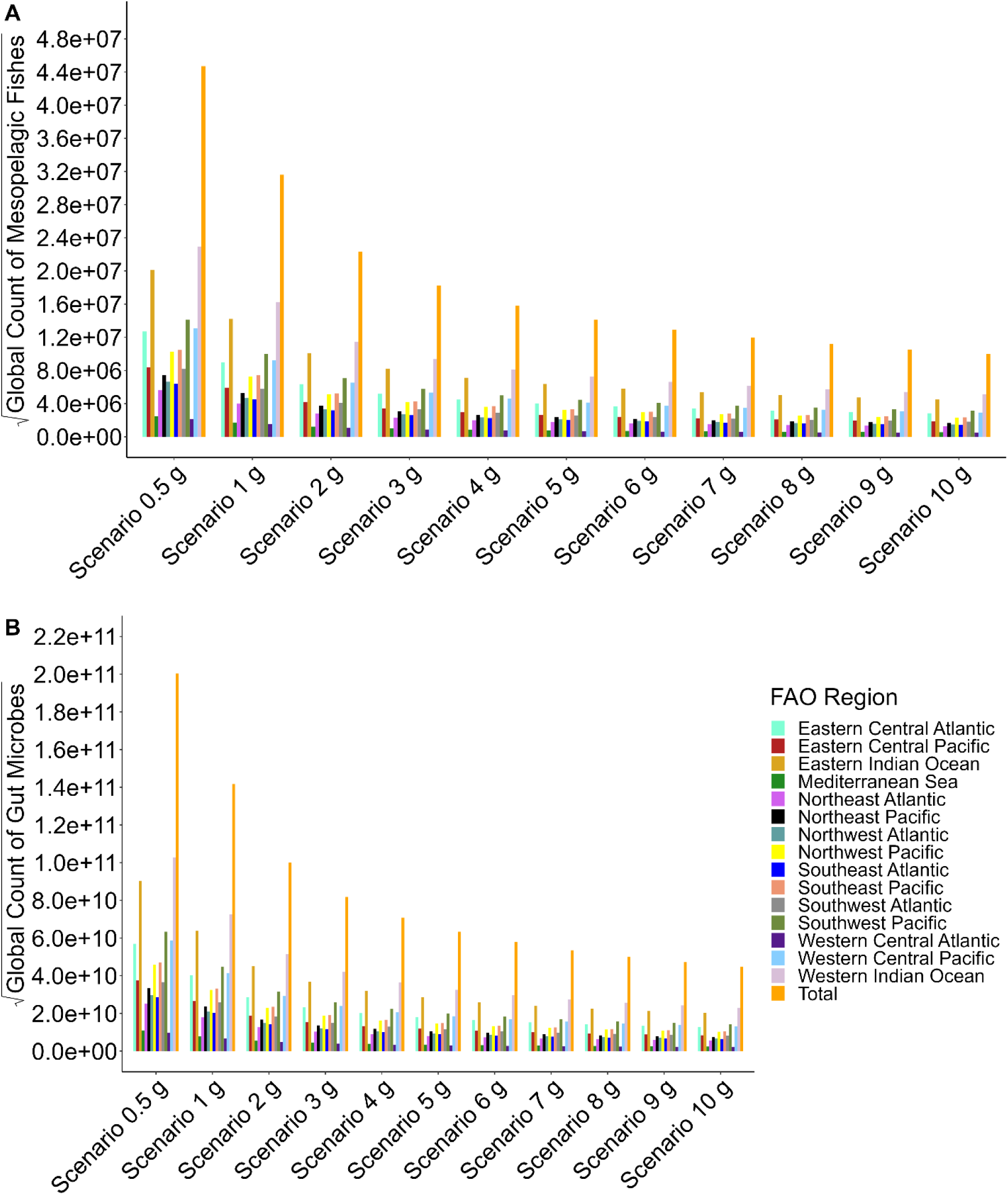
A) Global estimates of mesopelagic fishes aggregated and disaggregated by FAO region and B) global estimates of mesopelagic fish gut microbes aggregated and disaggregated by FAO region. Square root transformation is for graphical clarity. Note difference in scales. See Figure S1 for mapped biomass data.

**Table 2.**
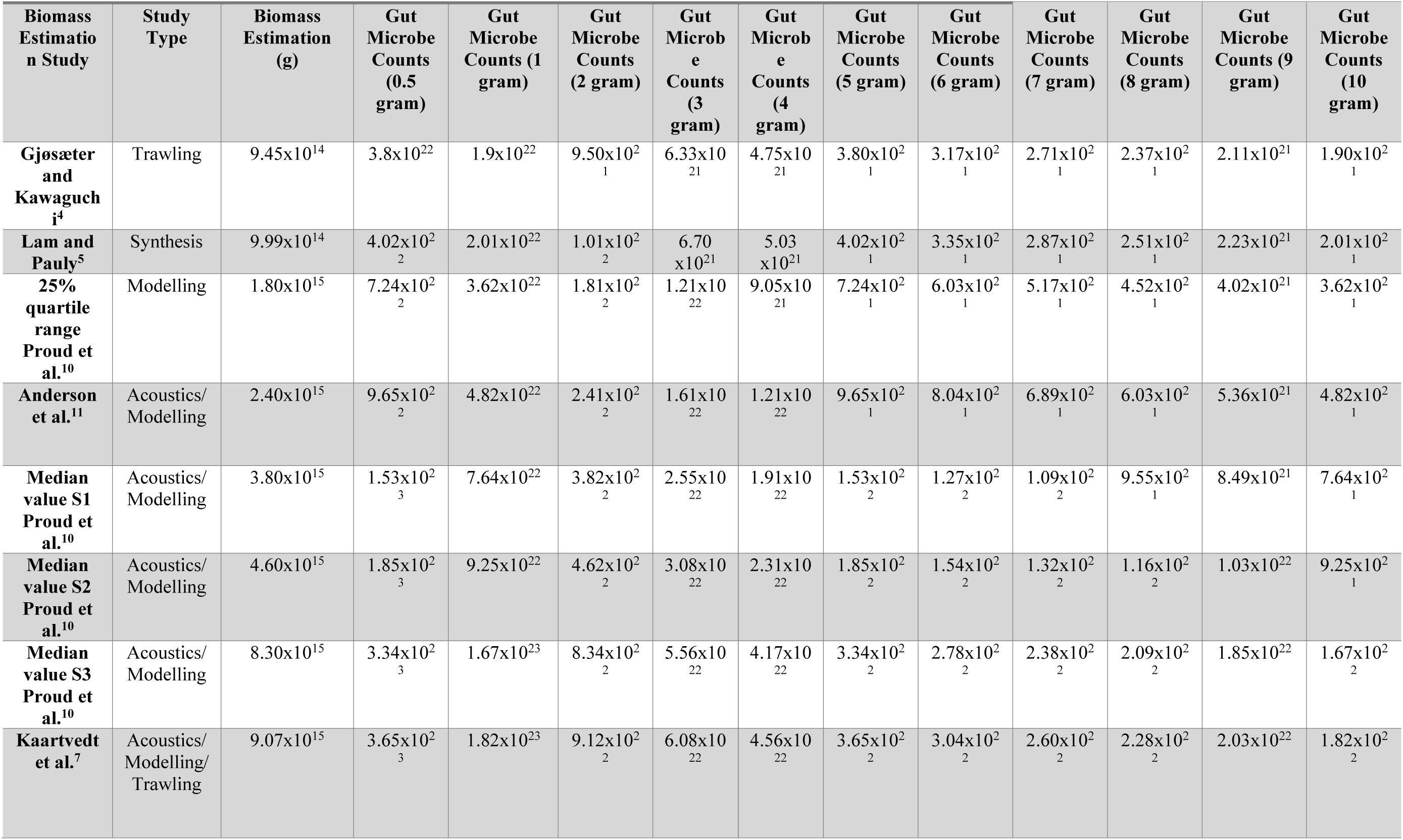
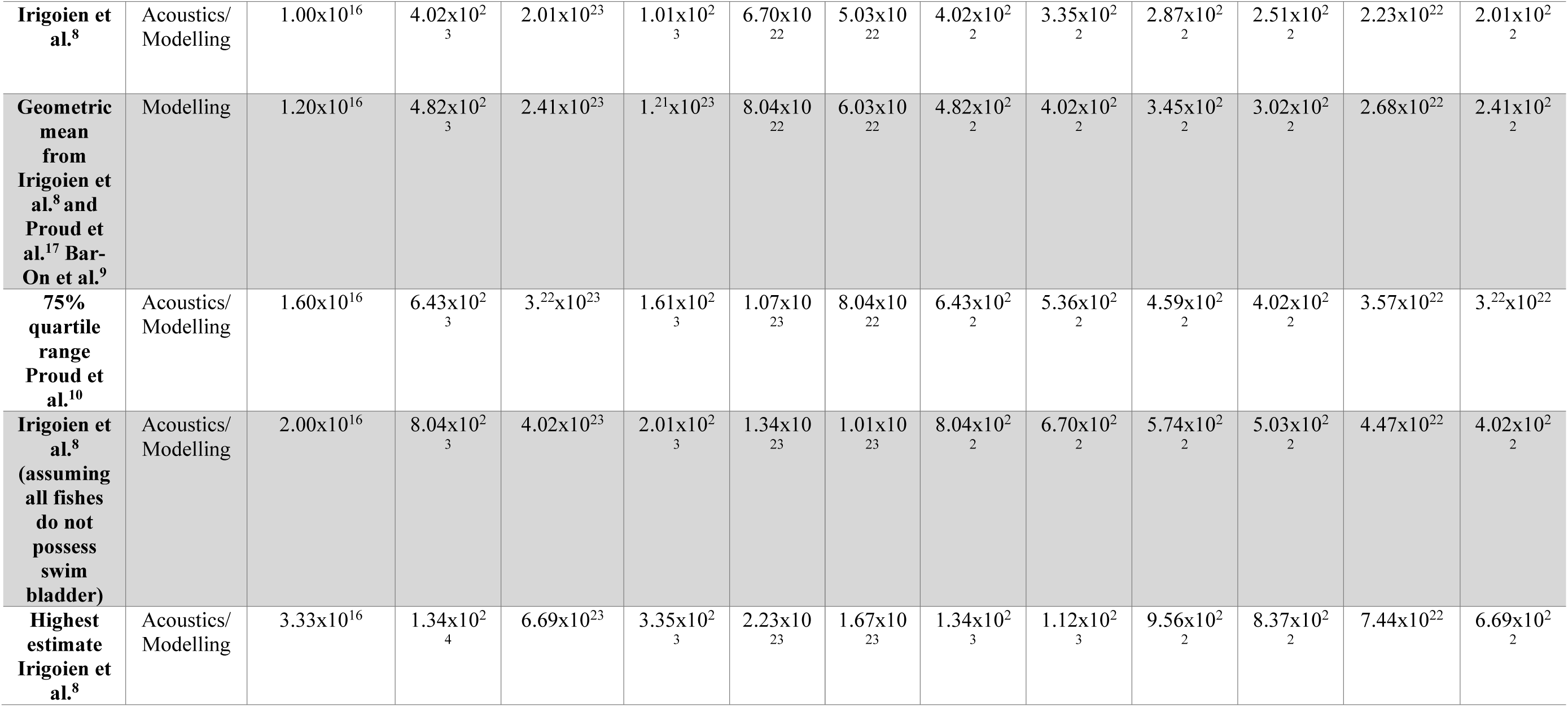
Global estimation of microbial load in mesopelagic fish gut microbiomes using the 11 individual fish mass scenarios (0.5-10 g) and acoustic, modelling, and trawl-based studies of total mesopelagic fish biomass.

### Metagenomics

A total of 24 frozen and 28 museum gut microbiome samples had metagenomic libraries prepared (Figure 1; Tables S1-S2). Samples for metagenomics are from different individuals than those used for flow cytometric analyses. Of these libraries, 24/24 (100%) of the frozen samples were created successfully, whereas 22/28 (79%) were successfully generated for museum samples. Whole-genome-shotgun sequencing yielded an average of ∼224 million paired-end reads per library and ∼120 million high-quality merged reads per library after preprocessing (Tables S2 and S3). Despite the higher DNA concentrations, quality, insert sizes, and sequence length distributions from frozen samples (Figure S2; Table S2), the lower biomass museum samples harbored more merged microbial reads (Figure S2; Table S2). Further data regarding preprocessing and fraction of aligned reads can be viewed in Table S2.

The taxonomic classification for prokaryote phylum, class, and family, as well as ordination of frozen and museum gut microbiomes using phylum-level taxonomic annotations is displayed in Figure 5. Phylum level taxonomic annotations yielded a statistical difference between deep-pelagic fish family (PERMANOVA, *p*=0.001, R^2^ = 0.116), preservation methods (PERMANOVA, *p*=0.001, R^2^ =0.174), and migration behavior (PERMANOVA, *p=*0.001, R^2^ = 0.091). While there was minimal taxonomic separation between migrators and non-migrators, this result was not statistically significant (ANOSIM, *p*=0.162, R=0.04). Pseudomonadota was the dominant phylum in 46 of 48 samples (Figure 5A). Similar reproducible taxonomic trends were observed for class and family level annotations, with genera from the class Gammaproteobacteria and family Moraxellaceae being generally most abundant across all samples (Figure 5BC).

**Figure 5.**
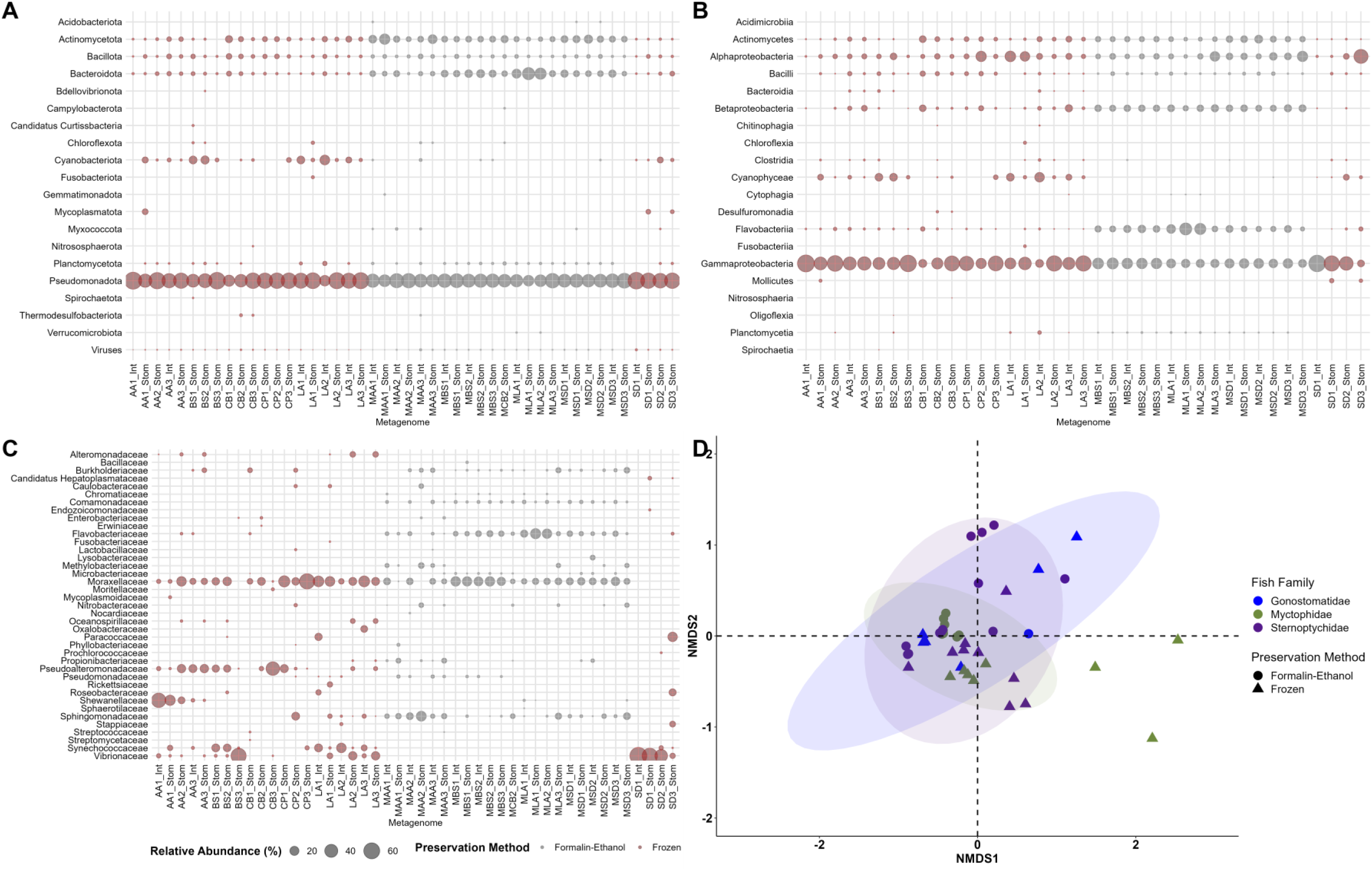
A-C) Phylum, class, and family level taxonomic annotations for mesopelagic fish gut microbiomes in frozen (red) and formalin-fixed (gray) specimens. D) Phylum-level taxonomic ordination of mesopelagic fish gut microbiomes by family and preservation method.

When analyzing the functional metabolic potential across samples, Amino Acid Transport and Metabolism was the most abundant COG Category in 39/45 (87%) of samples, accounting for >50% in most samples, followed by Coenzyme Transport and Metabolism and Replication Recombination, and Repair (Figure S3). Ordination of frozen and museum gut microbiomes using KEGG annotations is displayed in Figure S4. Gene catalogs can be viewed in Dataset S1. As with taxonomy, there was a statistical difference in functional metabolic potential between deep-pelagic fish family (PERMANOVA, *p*=0.001, R^2^=0.077), preservation methods (PERMANOVA, *p*=0.001, R^2^=0.104), and migration behavior (PERMANOVA, *p*=0.006, R^2^=0.046). While there was minimal functional separation between migrators and non-migrators, this result was not statistically significant (ANOSIM, *p*=0.801, R=-0.042). The reference-guided coassembly of all mesopelagic fish gut microbiome libraries revealed 96 complete and 36 nearly complete (>75%) pathways. Pathways of interest included those responsible for amino acid metabolism, cofactor and vitamin synthesis, and lipid metabolism, as well as ammonification and the urea cycle. Key activities and completed KEGG metabolic pathway completion for the coassembly can be viewed in Figure 6, and pathway completion for individual samples can be viewed in Dataset S2.

**Figure 6.**
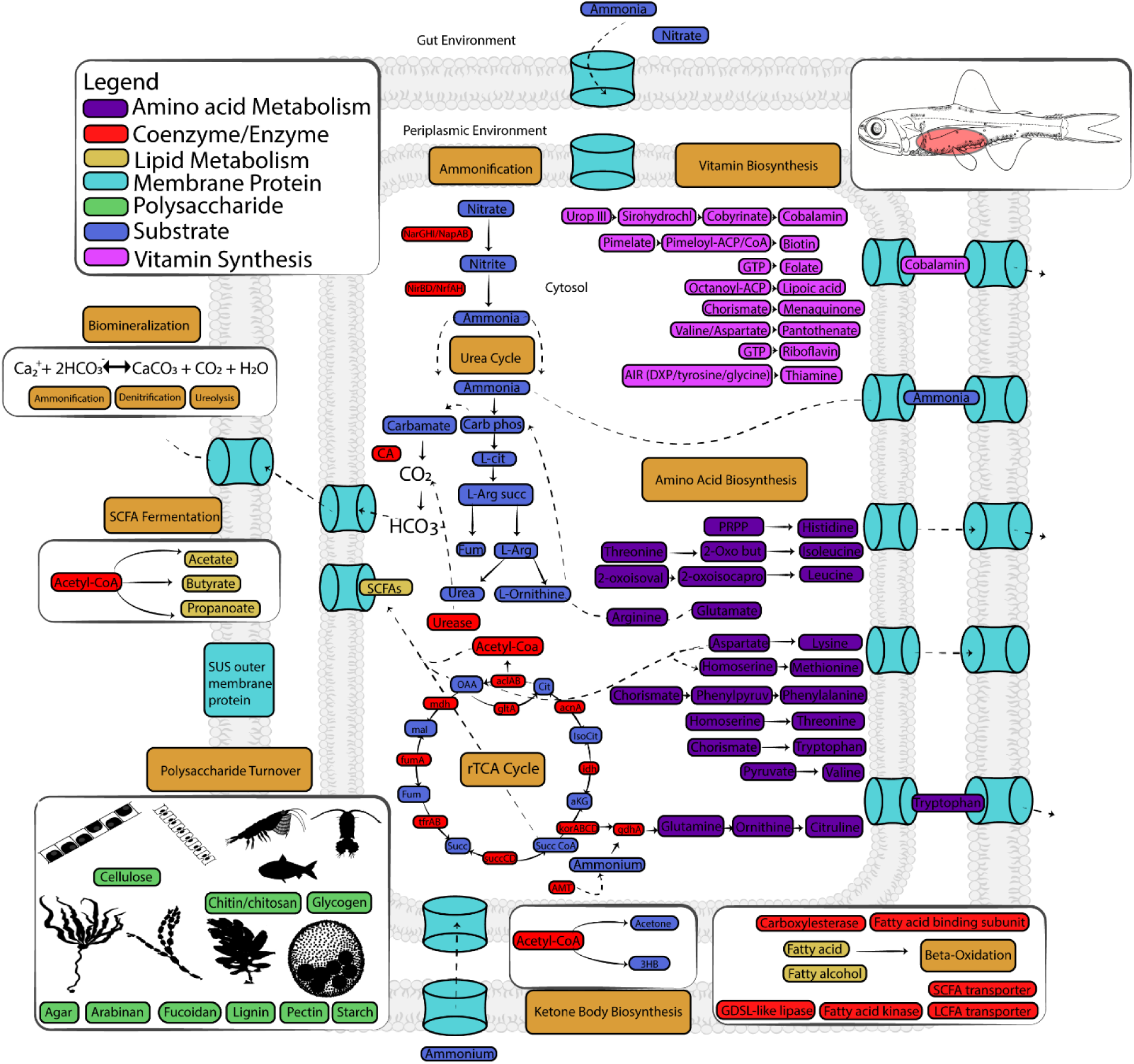
Key pathways and overarching functions of mesopelagic fish gut microbiomes. See Dataset S2 for full module completion data.

The clustering of polysaccharide degradation potential of mesopelagic fish gut microbiomes is displayed in Figure 7. Starch, glycogen, cellulose, chitin, and peptidoglycan degradation potential were the top five most common functions. The mean number of substrates that could be utilized by gut microbiomes across all samples was 7.59 ± 4.65, with a maximum of 19 possible compounds. See Dataset S1 for all CAZyme annotations.

**Figure 7.**
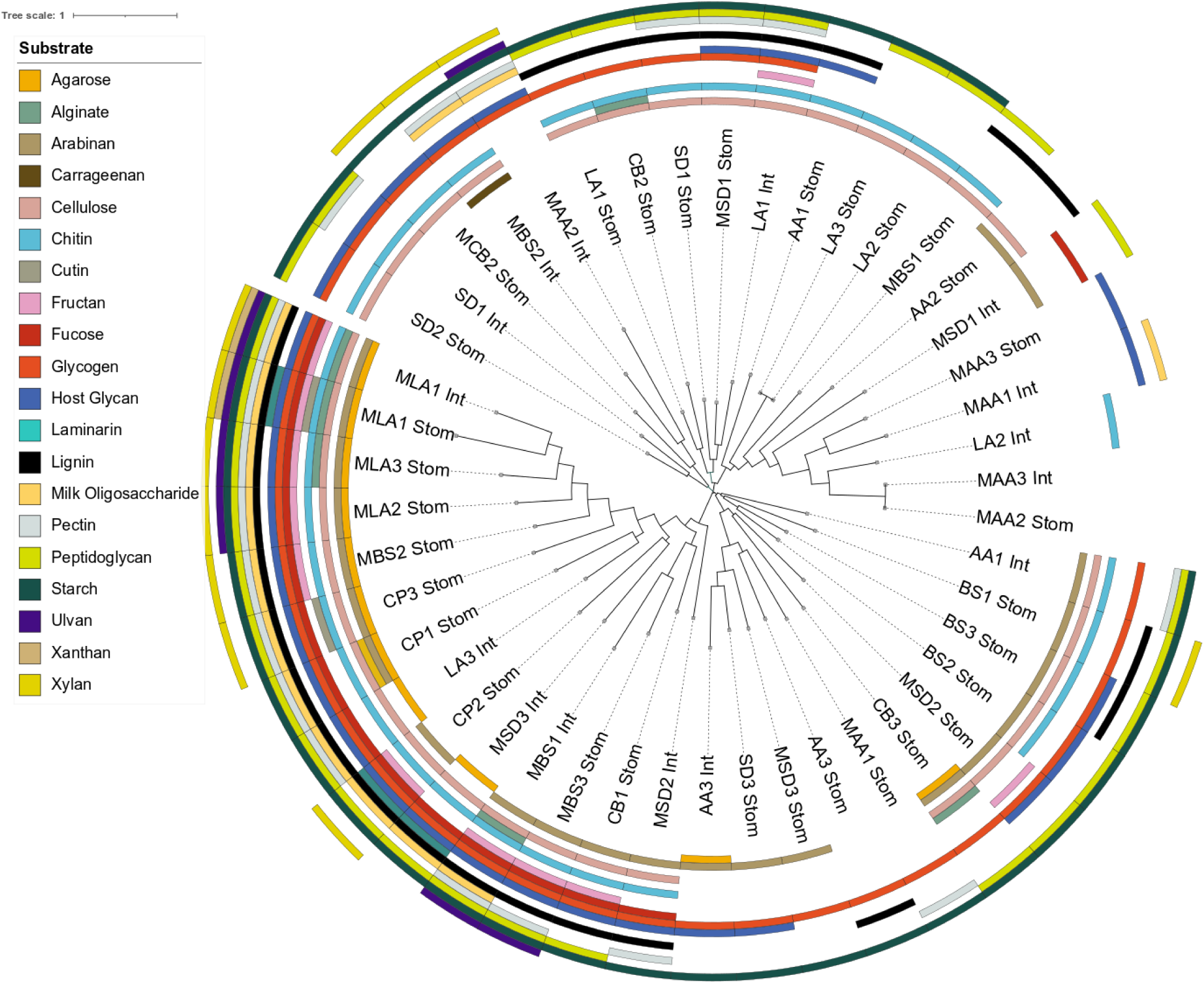
Animal and plant-based polysaccharide degradation potential of mesopelagic fish gut microbiomes. See Dataset S1 for functional annotations.

Two MAGs were binned and annotated from the coassembly, and the MAG taxonomic affiliations, as well as aligned functional annotations and open reading frames are displayed in Dataset S3. The environmental distribution of these MAGs is displayed in Figure S5. The *Acinetobacter junii* and *Psychrobacter* sp. MAGs harbored 43 and 47 completed or nearly completed metabolic pathways, respectively. Like for the coassembly, these MAGs harbored pathways for amino acid metabolism, cofactor and vitamin synthesis, and lipid metabolism, including the glyoxylate cycle (Dataset S2 and Dataset S3).

## Discussion

The traditionally embraced ‘holobiont’ perspective (i.e., analyzing fecal pellets, the end-product of host and microbial processes) to quantify oceanic nutrient flux has furthered our understanding of the biological- and microbial carbon pumps and carbon sequestration mechanisms^30,35,55,56^, nutrient remineralization patterns (e.g., pellet sinking rates and microbial activity on particles)^26–28,35,57,58^, and links between oceanic food webs and biogeochemical cycles^26–29,31–37^. However, this approach has limitations. Microbial transformations are often the bottleneck of biogeochemical processes, yet only knowing holobiont-level fluxes (e.g., fecal composition) does not enable us to understand how the environment and therefore trends in nutrient cycling impact and are impacted by gut microbes. Moreover, evolutionary dynamics of gut microbes are crucial for long-term biogeochemical cycling and animal health yet are hidden from analyzing fecal pellets. Additionally, gut microbes respond rapidly to environmental change (e.g., alteration of enzyme kinetics), whereas host metabolic potential is limited and physiology changes relatively slowly. The holobiont-only approach therefore misattributes all biogeochemical contributions to the host and “misses” biogeochemical hotspots over large spatial and temporal scales. We posit that our understanding of global biogeochemical frameworks is incomplete and has been limited by the absence of the data we present herein. These data markedly advance our understanding by yielding component-level insights on gut microbiome contributions that provide a mechanistic foundation for how these communities contribute to shaping biogeochemical and ecological processes in the water column.

### Mesopelagic fish gut microbiome estimates

This study provides the first estimates of global mesopelagic fishes (see supplemental), as well as approximations of the global mesopelagic fish gut microbial reservoir using published estimates of mesopelagic fish biomass^4–13^. While mesopelagic fish biomass is no doubt higher than 1 Gt (see supplemental)^4^, the estimates of gut-prokaryote abundance, and thus their potential impact, are still striking even at their absolute minimum. Altogether, the flow cytometric data collected from mesopelagic fish guts reveals an enormous microbial reservoir of 2.4×10^20^-1.34×10^24^ prokaryotes, though this range reflects substantial uncertainties in both fish biomass estimates and the representativeness of our microbial density measurements across all mesopelagic species and regions. We cautiously underestimate global prokaryote load in fish guts, as we only considered 1 ml of gut supernatant (5% of the available supernatant) for our estimates, and the volume of the digestive tract of many mesopelagic fishes exceeds 1 ml and may consequently disproportionately contribute to global gut microbial load. The most robust, current estimates by Kaartvedt et al.^7^ suggest that mesopelagic fish biomass may be 9 Gt, which when considered for deriving the total number of mesopelagic fishes, assuming an average mass of 3 g per fish, yields 6.08×10^22^ gut prokaryotes (Table 2; see supplemental for logical framing). Though the abundance of gut microbes in all mesopelagic fishes alone are less than the free-living load in the mesopelagic, their concentrations are two to three orders of magnitude higher than comparable volumes of seawater and are co-concentrated with organic matter. This begs the question of how heterotrophic production in global marine gut microbiomes compares to the surrounding water column (discussed below). With the inclusion of parallel contributions from highly abundant micronekton crustaceans (e.g., lophogastrids, mysids, and shrimps), which in many regions equal or exceed fish numbers, the abundance and biogeochemical contributions of gut microbes may begin to rival or exceed that of the free-living microbial reservoir.

The species of cosmopolitan mesopelagic fish families from various depths and ocean basins (and with different prey preferences and vertical migration behaviors^20,59–63^) harbored a range of 5.8×10^5^ – 7.8×10^7^ gut microbes ml^-1^. The observation that gut microbial counts were positively correlated with standard length, fish wet mass, and wet gut mass were largely consistent with trends predicted by an allometric trend scaling model^64^. We consider these gut microbial abundance estimates to be conservative and reasonable for the following reasons. First, the flow cytometric data presented in this study demonstrate similar normalized gut microbe abundances between five of the six frozen and museum collected species (Figure 3; Table 1). The total gut microbial counts ml^-1^ observed here are on par with some of the most highly abundant marine animals and terrestrial insects^64,65^. For example, the Antarctic Krill *Euphausia superba* is estimated to have a global biomass of 3.79×10^14^ g^66^ and have a range of 1.08×10^8^-2.51×10^9^ microbes g^-1^ ^66,67^ yielding about 1.9×10^23^ gut microbes^64^. Second, our estimates fall below the predicted microbial abundance of 3.1-6.9×10^24^ predicted to be found within all fishes on Earth (note that mesopelagic fishes likely account for 90% of global fish biomass and this was not included in the former calculations) and therefore below the estimated prokaryote abundance for terrestrial and marine animal-associated microbes, which was assessed at 1.3-1.4×10^25^ and 8.6-9.0×10^25^, respectively^64^.

The density of microbes per unit volume within animal guts is also one to three orders of magnitude higher than the surrounding seawater. Epi-, meso-, and bathypelagic waters have been shown across many studies to harbor a median free-living prokaryote abundance of 5.27×10^5^ cells ml^-1^, 1.51×10^5^ cells ml^-1^, and 0.43×10^5^ cells ml^-1^, respectively^54^. Oligotrophic, deeper waters (>200 m) can harbor 5×10^4^ cells ml^-1^ ^68^. In contrast, mesopelagic fish gut microbiomes contain a mean of 2.01×10^7^ ± 2.15×10^6^ SE microbes ml^-1^ and the observation that gut microbes ml^-1^ is statistically similar between migrators and non-migrators demonstrates that, even at depths where free-living prokaryotes decrease, gut microbiomes remain at least two to three orders of magnitude higher than surrounding seawaters. We posit that the gut microbiome of mesopelagic fishes are more dense and potentially more active because of greater abundances of organic and inorganic matter along the digestive tract that can stimulate metabolic activity and support growth. In support of this notion, one previous study of hatchetfishes and lanternfishes showed that the presence of organic material in the gut was often correlated with higher concentrations of viable microbes^50^. This supposition is further supported by a study of the gut microbiomes of deep-sea benthic organisms like abyssal amphipods and holothurians^69–71^, where the authors demonstrated that gut bacteria, which ranged in densities of 10^4^-10^9^ cfu g^-1^ responded to nutrient availability (e.g., host-ingested matter). Culturable cell counts have also been shown to be higher at *in-situ* pressure in larger deep-pelagic fishes^72^ and gut microbes may utilize organic matter more rapidly at depth^70,73^ suggesting increased activity relative to seawater communities.

As mentioned, our museum samples were fixed with 4% formalin and subsequently preserved in 70% ethanol. We hypothesized that fish cells would shrink due to dehydration as this has been seen in certain taxa where cells can shrink to ∼ 50% of their original volume in as little as 50 days^74–78^. As such we were concerned that host cells might be confused for microbial cells, but our calculation (shown below) and our flow cytometric method allay this concern. For example, we can calculate the diameter of the host cell (e.g., erythrocytes, immune cells). The mesopelagic fish *Valenciennellus tripunctulatus* possesses spheroidal erythrocytes ∼7 µm and *Maurolicus muelleri* has ellipsoidal erythrocytes ∼5-7 x 3µm^78,79^, the smallest host erythrocytes observed in teleost fish^68^, which usually have ranges of 8-16 µm x 6-11 µm in size^80,81^. Assuming a max of 50% volume shrinkage after long term ethanol storage, the spheroidal erythrocytes would have a diameter of roughly 5.56 µm and the ellipsoidal erythrocytes would have a volume of 11.78 µm^3^, which do not coincide with the lengths and volumes of bacterial standards nor cells gated in this study. Even if host cells picked up our stain, they would be larger than 3 µm and roughly quadruple the median volume of microbial cells after long term ethanol storage^81^ and therefore excluded from flow cytometric microbial counts.

### Gut microbiome metagenomics

There is a paucity of genomic studies characterizing gut microbiome composition of midwater animals and fishes^51–53^. Former studies suggest a low diversity of gut microbes but owing to the lack of taxonomic data, no core gut microbiome has been established. Our data reproducibly demonstrate, irrespective of preservation method, that there is a distinct type of microbial community either residing in or passing through mesopelagic fish guts (Figure 5). This observation is particularly striking considering that our samples range between 3-58 years old and come from different geographic regions (although Moraxellaceae members have been noted in modest abundances in other mesopelagic animal guts^53^). Broadly speaking, Moraxellaceae genera in fish gut microbiomes (i.e., *Acinetobacter* and *Psychrobacter*) make significant contributions to animal health through the fermentation of complex molecules into usable metabolites for the fish and synthesis of essential amino acids and vitamins to supplement nutrition^82^. Many of these pathways are observable in our MAGs (Dataset S3). As Moraxellaceae genera have been shown to be reproducibly abundant in animal-based diets^83^, their abundance in mesopelagic fish guts likely indicates protein fermentation by the gut microbiome, an energetically favorable microbial process^84^. Our *Acinetobacter junii* and *Psychrobacter* sp. MAGs have been observed predominantly in wastewater and seawater, respectively (Figure S5), yet appear to have gene sets suitable for symbiosis with host organisms, suggesting dual lifestyles to persist until reentering the gut environment (Dataset S3). *Psychrobacter* evolved from a host-associated lifestyle with mammals^85^, but the deep *Psychrobacter* ecotype harbors adaptive traits, which led to niche diversification in the deep-pelagial. Our *Psychrobacter* MAG appears to be psychro- and piezotolerant, with cold-adapted proteins and osmoregulatory mechanisms, as well as bile-salt symporters to withstand the pressures of the gut environment in deep waters (Dataset S3). Altogether our data suggests that *Psychrobacter* supplements mesopelagic fish nutrition and may continue to process fecal matter upon being excreted by the fish. *Psychrobacter* therefore may also control how colonizers of fecal pellets respond, but this remains to be determined.

There are multiple lines of evidence that these gut microbiomes could supplement host nutrition. In the coassembly, the observation that all essential amino acids and quintessential vitamins can be produced by the gut microbiome suggests that gut bacteria can complement or supplement host nutrition or may be required to provide these compounds. B-vitamins like cobalamin, niacin, and riboflavin have been shown to be limiting in fish diets^86^ yet are known to enhance growth and welfare^87^. Fermentation of amino acids yields short chain fatty acids such as acetate, butyrate, and propionate, which can serve as energy stores for the host as well as protect gut barrier function^88^. Similarly, ketone bodies, which are energy-rich molecules derived from fatty acid biosynthesis, can reduce gut inflammation resultant from lipid-rich diets and be utilized by the host during periods of food scarcity^89^. As such the gut microbiome may contribute to some organisms’ ability to subsist for long time periods between successful predation events. Metabolites produced by the gut microbiome such as SCFAs and ketone bodies may reduce the metabolic investment to vertically migrate, or upon being sated, GABA production by the gut microbiome may promote a relaxed state^90^ and prevent migration. Altogether, the gut microbiome plays an important role in modulating animal health, which in turn shapes the prevailing biogeochemical and ecological processes over large spatial and temporal scales in the pelagic water column.

The gut microbiome may also increase the transfer efficiency of organic matter that would otherwise “escape” host digestion and be cycled in surrounding seawaters. Myctophids forage on energy rich prey (predominantly copepods) to compensate for decreased efficiency and may not utilize ingested material to the fullest extent. The current body of evidence suggests that migratory myctophids have a high throughput, lower efficiency digestive tract^91^, with short guts (relative to their body size), lesser nutrient absorption^92^ and lower enzymatic activities^93^. Proteobacterial genera are dominant in mesopelagic fish guts^51–53^ (Figure 5), but this is distinctly different from mammals, which are dominated by Firmicutes and Bacteroidetes that have been shown to be less efficient at degrading gut derived carbohydrates^94^. These Proteobacteria in mesopelagic fish guts likely play a distinct role in global oceanic polysaccharide turnover. Indeed, the gut microbiome harbors the potential to degrade abundant polysaccharides such as cellulose and chitin, among many other animal and plant-derived compounds (Figure 7). Cellulose is the most abundant biopolymer on Earth and is the main constituent of algal and plant cells.^95^ Therefore, gut microbiomes are likely to expand fish metabolic capabilities, as many mesopelagic fishes are not thought to be capable of degrading cellulose, lacking the necessary enzymes to do so. Chitin, a major component of marine exoskeletons, is a known biodegradable compound that has existed for millions of years and annual chitin production is approximately 10^10^–10^11^ tons^96^. In a separate study, chitinase activity of three myctophid fish species was measured by pooling four to five individuals and adding Β-glucosidase to measure N-Acetyl Glucosamine production^93^. The authors of that study demonstrated that myctophids have high chitinase activity and attributed this activity to digestive morphology, with the intestinal tissues (including organic matter) exhibiting chitinase activity increases up to 600% after the addition of Β-glucosidase. While the authors concluded that microbial activity could not be ruled out, chitinase activity was largely attributed to the hosts, although microbial contributions remain to be studied. Microbial contributions, in addition to animal attributes (e.g., digestive morphology, feeding periodicity, gut pH, gut transit time), could explain how mesopelagic fishes like *Ceratoscopelus warmingii* may derive energy from ingested algal material^97^.

The contribution of gut microbiomes to nitrogen cycling have yet to be considered in nitrogen flux models, but free-living microbes make significant contributions to the oceanic nitrogen cycle^98^. As our data suggest, deamination of organic nitrogen compounds (e.g., chitosan production, protein decomposition) is common among mesopelagic fish gut microbiomes, and the ammonification and ureolysis pathways are well-represented, so midwater gut microbiomes are likely significant drivers of nitrogenous compound production (e.g., NH_3_, NH_4_^+^, N_2_O, NO_2_, NO_3_^-^). While excretion by fishes influences the distribution of nitrogenous wastes in shallow and deep-pelagic waters, microbial contributions to ammonia production likely directly influence the extent of gut nitrogen production and how the surrounding water column responds (e.g., phytoplankton growth). Similarly, fishes contribute substantially to the marine inorganic carbon cycle through insoluble calcium carbonate (CaCO_3_) production in the gut^6,99,100^ as an osmoregulatory byproduct of drinking seawater. However, the role gut microbes may play in this biomineralization pathway in mesopelagic fishes has not been explored, and this precipitation has been attributed entirely to fish hosts^9^. Polymer substances like abundant cellulose and chitin (potential template for carbonate production) may stimulate heterotrophic metabolism and enable microbial precipitation of CaCO_3_ via genes (i.e., carbonic anhydrases) and gene sets for well-represented pathways in the gut such as ammonification^101^, denitrification^102^, and ureolysis^103^.

### Considerations for the biological and microbial carbon pump concepts

Mesopelagic fish gut microbes alone are within four orders of magnitude of the entirety of the free-living prokaryote reservoir at the same depths, and with the inclusion of the rest of the biomass at mesopelagic depths the disparity likely decreases. This striking abundance of gut microbes raises the question as to their net contribution to marine biogeochemical cycles. In addition to being a significant fraction of the mesopelagic microbial community, gut microbes that have a differentially larger impact via two other factors. First, the gut microbes of migratory animals likely have broader spatial impact than bacterioplankton as they are presumably metabolically active throughout the migration, transforming organic matter along the way. Migratory fish gut microbiomes likely add a spatial component to the microbial carbon pump concept^56,104^ whereby microbial denizens of the gut, in synergy with host digestive enzymes, transform organic matter between labile and recalcitrant states. Midwater animal gut microbiomes may therefore be important sources of carboxyl-rich alicyclic molecules (CRAM)^105^ or the eventual breakdown of CRAM analogs in the pelagic water column and should remain a topic of future metabolomic studies. The migration of the host fishes likely expedites the flux of these compounds to depth and alter the fate of carbon. Second, gut microbes may well be more metabolically active because their host fish acquires and concentrates organic matter via feeding. This second point remains to be tested, but it is highly plausible given data from other free-living, particle-associated, and host microbiome studies^50,57,58,68–71^. Intuitively, gut microbiomes may therefore be microbial seed banks for the water column, accounting for a significant fraction of the 3.026×10^28^ prokaryotes in mesopelagic seawaters and may alter how free-living microbes respond to fecal matter. Understanding what fraction of gut microbes survive transit through the digestive system and remain active post-excretion is a logical area of inquiry. These gut microbiomes, alongside free-living organisms, likely have further synergies and enable successive processing of matter in the water column, which will have different degrees of impact on the biological and microbial carbon pumps.

### Museum samples as viable tools to study the past

The formalin-fixed museum samples (13-58 years old) outperformed the frozen samples (3-8 years old) of lesser age in terms of percent microbial sequences present out of total raw reads (0.83% vs. 0.06%), though of lesser quality (Figure S5). This is enigmatic considering the frozen libraries were sequenced more deeply and were of better quality (Figure S5; Table S2), requiring no extra processing to remove dimers unlike for the formalin samples. We hypothesize that the disparity in microbial sequences arises from either freeze/thaw/lysis or subsequent agitation and heating (to assist with proteinase K digest) of bacterial cells in the frozen samples and not implementing a dedicated nucleic acid fixation step. While utilization of frozen, unfixed samples has revealed microbiome taxonomic differences between samples at shorter timescales (8 weeks)^106^, and our data suggest this can still be the case after 3-8 years, these libraries are fragmented (in relative percentage of microbial sequences) likely because of destruction of microbial DNA from prolonged heating. Answers may be found in the flow cytometry workflow; for reproducibility in microbial counts, frozen samples were fixed in formalin before processing and had statistically similar gut microbes ml^-1^ to museum samples, indicating that the bacteria are present. This observation suggests that prolonged heating at 60 ℃ may have facilitated heat-induced destruction of microbial DNA. Future studies working with aged frozen samples should stabilize DNA while thawing or prior to freezing. Importantly, while similar conclusions have been drawn regarding the functional metabolic potential of midwater fish gut microbiomes (discussed in previous paragraphs)^51–53^, we have arrived at similar conclusions using aged frozen and museum samples.

### Conclusions

Mesopelagic fish gut microbes have been an unconstrained component of marine mesopelagic biogeochemical cycling. These data illustrate that they are sufficiently abundant to warrant consideration when thinking about biogeochemical processes, and the likelihood that they are more active than their free-living counterparts further raises the possibility that they may have a marked impact in shaping global oceanic nutrient cycling. This being said, future studies should focus on establishing mesopelagic fish gut microbe metabolic rates and work to determine their contributions to the respective processes. Moreover, the survival and continued activity of gut microbes after excretion remains unclear, which affects their potential contribution to water-column processes. Establishing the baseline patterns and processes of global midwater gut microbiomes, including how microbial biomass, gut microbiome functions, and metabolite concentrations present in digesta change across the mesopelagic realm in other biomass abundant animals, and -ultimately-quantifying microbial contributions to carbon flux and sequestration on regional and global scales are necessary steps. Calculating ocean energy turnover by gut microbes and incorporating these data into biogeochemical models may enable us to establish a more comprehensive appreciation for global oceanic nutrient cycles that govern the very conditions of our biosphere.

## Methods

Archived mesopelagic fish samples were retrieved from the MCZ at Harvard University. The ecologically relevant species chosen for this study were based on known abundance and biomass data and are considered to be ecologically representative at the family level (e.g., Gonostomatidae, Myctophidae, Sternoptychidae, Stomiidae). Frozen mesopelagic fishes were collected onboard the RV *Point Sur* in the Gulf of Mexico as part of Deep Pelagic Nekton Dynamics of the Gulf of Mexico (DEEPEND) Consortium research. These fish species were classified as being migratory or non-migratory using previous studies^20,59–63,107^. The locations where samples were originally collected from are displayed in Figure 1. Sampled metadata are provided in Table S1.

Fishes were removed from sample jars using sterile, precombusted forceps, and placed into sterile petri dishes. The exterior of the animal was generously rinsed with 70% ethanol to remove potential adhered cells and organic material. Standard lengths of fishes were measured as the distance from the tip of the rostrum to the end of the hypural plate^108^. The wet mass of the fish and wet mass of the gut were measured using gravimetric technique. Dissections were performed using precombusted forceps and surgical blades in petri dishes. Dissection tools were cleaned with ethanol and flame in between samples. An incision was made spanning from the isthmus (area between gills on ventral portion of the body) to the insertion of the anal fin and the stomach and intestine were removed from the body cavity. The digestive tract (stomach and intestines) of each sample was opened with a longitudinal slice, chopped into smaller pieces and placed into vials containing filter sterilized (0.2 micron) and autoclaved phosphate buffered saline (PBS; 1x) and precombusted ceramic beads (2 mm). Potential microbial cells in the esophageal tissues and pyloric caeca were not included in this analysis. Gut suspensions were created by vortexing samples using previous methods^69,71^. Particulate material was allowed to settle for one hour (e.g., removal of larger host cells) and 1 ml of supernatant was transferred to an Eppendorf tube, subsequently split into two 500 µl fractions (unstained control and fluorophore positive), centrifuged for 1 minute at 7000 rcf to facilitate disaggregation of potential cell clumps^109^ and stained using 500X Bactoview Live Red^TM^ (Biotium, Fremont, CA; 1:500 v/v) for flow cytometry. Bactoview Live Red^TM^ has an excitation and emission of 572 nm and 675 nm, respectively, and is a cell permeable, DNA stain, with weak mitochondrial staining and a proclivity to stain dead cells with greater fluorescent intensity. We have investigated other potential fluorophores to quantify gut bacteria. While other membrane-embedding fluorophores (e.g., cardiolipin-specific), which also have weak mitochondrial staining are available, the staining efficiency of lipid-activated dyes are sensitive to the amount of lipid crosslinking.

### Flow Cytometry

Flow cytometry was conducted using an Aurora™ (Cytek Biosciences, Fremont, CA) at the Bauer Core Facility at Harvard University. On the Aurora™, 150 µl of both unstained and stained supernatant were aliquoted into U-bottom 96-well plates, with ∼30 µl dead volume and purging and SIT flush in between wells. After voltage optimization, a threshold of 500 was chosen to image small cells, and these cells were measured at low flow rate (15 µl/min) for a maximum time of 90 seconds, specifically using the yellow-green lasers (YG1-YG10). Data were processed in FCS Express (https://denovosoftware.com/) using a hierarchical gating strategy. Unstained controls for each fish individual were used as a guide to determine fluorophore positive populations in fish guts. Using density plots, stained cells were gated by including densities (counts) greater than 1 to refrain from including potential doublets in downstream analyses. Next, the log transformed FSC-A and SSC-A of stained cells representative of populations isolated from the gut were extracted and subsequently plotted against one another. Control beads and bacterial standards with operationally defined scattering information and size were overlayed on scatter plots of stained cells. Stained cells less than 3 microns were gated in the same manner as described above to exclude any potential doublets.

Control bacterial standards *Corynebacterium xerosis* (gram positive), *Shewanella oneidensis* (gram negative), and *Vibrio fischeri* (gram negative) were purchased from Carolina Biological Supply Co. (Burlington, NC). These bacteria were isolated from culture, fixed with 4% neutral-buffered formalin for 24 hours, washed with PBS twice, and stained with Bactoview Live Red^TM^ to provide objective gating based on the scattering characteristics of known bacteria and their sizes. In addition to using bacterial standards as a guide for gating, control polystyrene beads (Spheros) ranging from 0.2 µm – 7.4 µm (0.2 µm, 0.5 µm, 1 µm, 3.4 µm, 5.1 µm, 7.4 µm) were also used to inform gating efforts. Unstained and stained controls with the autoclaved, filtered-sterilized (0.2 µm) PBS were also measured to prevent contamination and establish potential background noise that could be confused with cells.

### Epifluorescence Microscopy

Stained supernatant returned from the Aurora™ was placed into a syringe and filtered through stacked polycarbonate (0.2 micron) and glass microfiber (0.7 micron) filters in a filtration apparatus. In the apparatus, the filter was washed with 1x PBS followed by two rinses of ultrapure water and then subsequently dehydrated with 50% and 80% ethanol for 30 seconds each. Filters were placed in darkness in a laminar flow hood to dry and successively cut in half using a sterile razor blade. Filters were arranged and mounted on glass slides with Vectashield ® (Vector Laboratories, Newark, CA) and stored at -20 ℃ for one hour before imaging using epifluorescence microscopy.

#### Global Count of Mesopelagic Fishes and Gut Microbes

Mesopelagic fish biomass estimates (g m^-2^) from Gjøsæter and Kawaguchi^4^ were leveraged to extrapolate gut microbes ml^-1^ to regional and global scales using shape files created by Lam and Pauly^5^ and ArcMap (ESRI). The clip tool was used to separate areas where shapes overlapped for more precise estimates of total area. Eleven scenarios (0.5 g, 1 g, 2 g, 3 g, 4 g, 5 g, 6 g, 7 g, 8 g, 9 g, 10 g; essentially one order of magnitude) of the average wet mass of mesopelagic fishes were used to estimate the number of fishes for each FAO region by using the biomass estimates from Lam and Pauly^5^, total number of mesopelagic fishes, and multiplying by the geometric mean 2.01×10^7^ microbes ml^-1^. As recent acoustic and modelling studies provide evidence that mesopelagic fish biomass is either equal to 1 Gt or higher^4–12^, total mesopelagic fish biomass data from these studies were used to estimate mesopelagic gut microbial load on a global scale using the same methods as described above.

## Metagenomics

### DNA Extraction and Library Preparation

Museum samples from the Harvard MCZ were classified as “museum” samples, whereas frozen samples collected from the Gulf of Mexico were classified as “frozen” samples (collection details are described earlier). Museum samples were comprised of fishes that were initially fixed in formalin for an indeterminate amount of time then transferred to 70% ethanol for long-term storage. All fishes were between 2 and 59 years old. Museum and frozen fish guts were excised as described above for flow cytometric analyses. With 25 mg input material, DNA was extracted using the QIAamp DNA Mini kit (Qiagen Sciences, Germantown, MD) following manufacturer protocols, with slight modification to digestion and elution steps. Specifically, Proteinase K digest was achieved overnight while heating at 60 ℃ at 300 rpm to fully digest tissues and multiple heated elution steps at 60 ℃ were used to unbind stubborn DNA from columns to increase yields. DNA concentrations were first assessed with Qubit™ fluorometry (Thermo Fisher Scientific, Waltham, MA) and then TapeStation™ model 4200 (Agilent Technologies, Inc., Santa Clara, CA), with Tapestation providing estimates of DNA fragment sizes. Frozen samples were library prepped using Illumina™ DNA ¼ volume kit (Illumina, San Diego, CA), whereas formalin-fixed samples were prepared using Kapa Hyper Plus ¼ volume kit (Roche Diagnostics Corporation, Indianapolis, IN). Libraries were randomly spot checked a second time to estimate DNA fragment sizes and pooled for sequencing. The formalin-fixed sample pool required removal of dimers, and dimers were removed using two 0.9x magnetic bead cleanups on that pool. Frozen and museum samples were sequenced separately to avoid preferential sequencing of smaller fragments and across two lanes on a NovaSeq™ X 25b at the Bauer Core Facility at Harvard University.

### Bioinformatic Workflow

Paired-end read libraries were first error corrected using tadpole.sh in error correction mode (pincer=t, tail=t) and subsequently merged with bbmerge.sh using default parameters and specifying adapter sequences^110^. Merged read libraries were quality trimmed using bbduk.sh (trimpolygright=6, minlen=70, hdist=2). The quality of merged read libraries was assessed using FastQC^111^. All read libraries were mapped to assembled host reads with bowtie2^112^ (-D 20 -R 3 -N 0 -L 20 -i S,1,0.50) for either the same species or closely related species to deplete host contamination. Mapping statistics can be viewed in Table S2. Host read libraries (genome skims) were downloaded from SRA using fasterq-dump, preprocessed using fastp^113^ with default parameters, and assembled using MEGAHIT^114^ (-t 64 -m 950 --presets meta-sensitive). MEGAHIT^114^ was used because it could be expected that a eukaryote genome would be complex and more than 2 TB memory was required for genome assembly with SPAdes^115^. Host depleted libraries were mapped to an index of nt_prok split into four chunks using bowtie2^112^ with the former parameters to recruit prokaryotic reads. Mapped reads were converted into .fa format using reformat.sh under the BBmap^110^ suite and taxonomically identified using blastn^116^ against nt_prok (e-value 1e^-10^, bitscore 100), as well as Kaiju^117^ (-E 1e^-4^; -s 65; -l 11; -e 3). Unique prokaryotic reads meeting the former criteria were pulled from the original host-depleted library (using sequence identifiers) and reformatted to .fa for creation of contig databases in Anvi’o^118^ Briefly, open reading frames of putative gene fragment sequences were predicted using prodigal and subsequently annotated using CAZymes^119^ (dbCAN2), COGs^120^, KOfam^121^, Pfam^122^ within the Anvi’o platform and subsequently catalogued with scripts available at https://github.com/Echiostoma/Mesopelagic-Fish-Gut-Microbiomes. KEGG module completion was determined using the anvi-estimate-metabolism command. Prokaryotic read libraries from all samples were concatenated and coassembled using MEGAHIT as described above. While a coassembly with metaSPAdes produced a larger total length, the N50 was smaller than that of MEGAHIT, and a majority of contig sizes were smaller than 1500 bp, so the coassembly with MEGAHIT was used for binning of genomes. Depth of coverage of mapped reads to the coassembly were assessed with runMetabat.sh and metagenome-assembled genomes (MAGs) were binned using Metabat2^123^, with a minimum contig size of 1500 bp. Marker gene completion and contamination for MAGs was determined using CheckM^124^. Manual refinement to remove duplicate copies of marker genes was done using the anvi-refine command. MAG taxonomy was determined using GTDB-Tk2^125^ (v2.4.0, release 220). MAGs were functionally annotated as described above.

To assess the ecological distribution of our MAGs, MAGs were submitted to the Branchwater Metagenome Query tool (https://branchwater.jgi.doe.gov/). This tool is an extension of the Sourmash^126^ platform (k-mer based taxonomy) that queries public metagenomes (>1 million metagenomes) on NCBI Sequence Read Archive. Matches that had a containment average nucleotide identity >0.97 were considered to be species-level matches, as suggested by the developers.

## Statistics

For flow cytometry, in-software gate statistics were used to calculate the cells encompassed by each gate and fluorescent intensity of the gated population per sample. The standard length (mm), wet mass (g), wet mass of the gut (g), and cell count data were non-parametric after applying several transformations. Therefore, non-parametric statistical analyses were applied throughout. To correlate standard length (mm), wet mass (g), and wet mass of the gut (g) with cell counts ml^-1^, Spearman’s Rank Correlation was used. A Mann-Whitney Wilcoxon test was used to statistically compare the mean cell counts ml^-1^ based on migration behavior (migratory, non-migratory). The Kruskal-Wallis test, with post hoc multiple comparisons was used to compare cell counts ml^-1^ between deep-pelagic fish families (Gonostomatidae, Myctophidae, Sternoptychidae, Stomiidae). Mann-Whitney tests were used to statistically compare normalized microbes ml^-1^ g^-1^ between frozen and museum specimens. All statistical tests were performed using R^127^.

## Acknowledgements

We thank Andy Williston and Meaghan Sorce at the Harvard MCZ Ichthyology collection for assistance with sample acquisition. We acknowledge Virginia Armbrust and Francois Ribalet for helpful discussions regarding flow cytometry. We thank Federica Calabrese and Ian Hughes for assistance with sample processing. We thank the Harvard University Bauer Core Facility for assistance with flow cytometry and library preparation and whole-genome-shotgun sequencing of samples. We would also like to thank the DEEPEND Consortium (www.deependconsortium.org) for sample collections.

## Data Availability Statement

Raw sequencing data to be deposited and embargoed at the NCBI Sequence Read Archive and made available after publication. Code used in bioinformatic workflow can be found at https://github.com/Echiostoma/Mesopelagic-Fish-Gut-Microbiomes.

