## Supplemental Information for "Microbial Communities in Mesopelagic Fish Guts Suggest an Overlooked Component of Marine Biogeochemical Cycles"

**This PDF file includes:**
Supplemental discussion
Figures S1 to S5
Captions for Tables S1 to S2
References for SI reference citations

**Other supplementary materials for this manuscript include the following:**

FCS files used for flow cytometric analyses
Code used for analyses
Datasets S1 to S3

### Supplemental Discussion

*Mesopelagic fish estimates*

We began this treatment with the caveat that while it is accepted that global mesopelagic fish biomass is quite high[1-8], possibly orders of magnitude higher than all global fisheries landings combined, a properly resolved estimate is one of the outstanding issues in biological oceanography. Working towards resolving this issue, in this study, mesopelagic fish biomass measurements[1-6,9-11] paired with potential average individual mesopelagic fish mass scenarios (0.5 – 10 g) revealed a first order estimate of $1.0 \times 10^{14}$ – $6.66 \times 10^{16}$ global mesopelagic fish individuals. While all mesopelagic fishes exhibit variable catchability in different gears due to size and mobility[12] and thus are not fully represented in global biomass data[1,8], the vast majority of mesopelagic fishes range from 2-15 cm total length[13]. Therefore, the range of masses for individuals of each species is generally constrained by the above size distribution. Bayesian model estimates of length-weight relationships of mesopelagic fishes could be used to predict the average mass of mesopelagic fishes on a species-level basis for the many mesopelagic fish species where no empirically derived length-weight regressions exist. That said, we posit that the average wet mass of mesopelagic fishes is significantly right-skewed (most values clustered towards the lower end) because of the numerical dominance of small fishes (e.g., *Cyclothone*, *Vinciguerria*)[8,14]. Indeed, while only a fraction of the ocean has been trawled, *Cyclothone*, thought to be the most abundant vertebrate, has been shown to constitute a significant portion of collected trawl biomass, and average *Cyclothone* length is roughly 3 cm. While included in this study, an average wet mass of 0.5 g per mesopelagic fish, that may very well apply to *Cyclothone* and *Vinciguerria* globally, is likely too small to apply to all mesopelagic fishes, whereas 10 g is potentially too high because again, the numerical dominance of small fishes significantly skews these data. Consequently, it is reasonable to hypothesize that an average wet mass between 3-5 g could be representative of mesopelagic fish assemblages. When paired with mesopelagic fish biomass estimates, we can use this wet mass estimation to crudely estimate the number of mesopelagic fishes. The highest modelled biomass estimates from acoustic backscattering data of 22 Gt [2] (assumes most fishes do not possess swim bladders) and 33 Gt [2] are likely overestimates of global mesopelagic fish biomass (Table 2). The estimate of 10 Gt [2] is also an unlikely representation, as this value attributes all backscatter to fishes and does not account for gas filled bladders of other animals (e.g., siphonophores). The acoustically determined estimate of ~9 Gt by Kaartvedt et al.[1], however, is a more realistic global figure of mesopelagic fish biomass[15]. If an average mass of mesopelagic fish ranges between 3-5 g and we pair these masses with the ~9 Gt estimate, there are $1.51 \times 10^{15}$-$3.02 \times 10^{15}$ fishes. While the global mesopelagic fish biomass is still open for study, at the very least, there is 1 Gt of mesopelagic fishes distributed globally[9], which yields $1.0 \times 10^{14}$ to $2 \times 10^{15}$ fishes assuming 0.5g and 10 g mesopelagic fishes, respectively. The number of fishes is within an order of magnitude (1 Gt vs. 9 Gt).

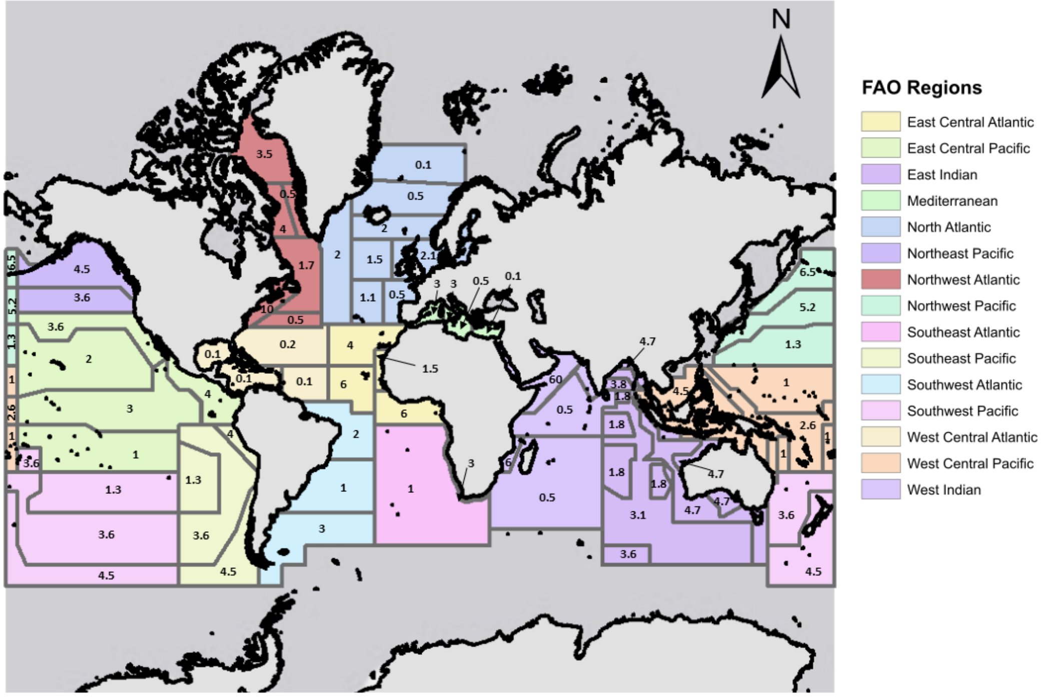


**Figure S1.** Mesopelagic fish biomass estimates (g $m^{-2}$) synthesized by Gjøsæter and Kawaguchi[9] from trawl data (~1 Gt total biomass). Figure adapted from Lam and Pauly[10]. Values displayed on shapes represent mesopelagic fish biomass (g $m^{-2}$). Colors of shapes represent defined FAO regions.

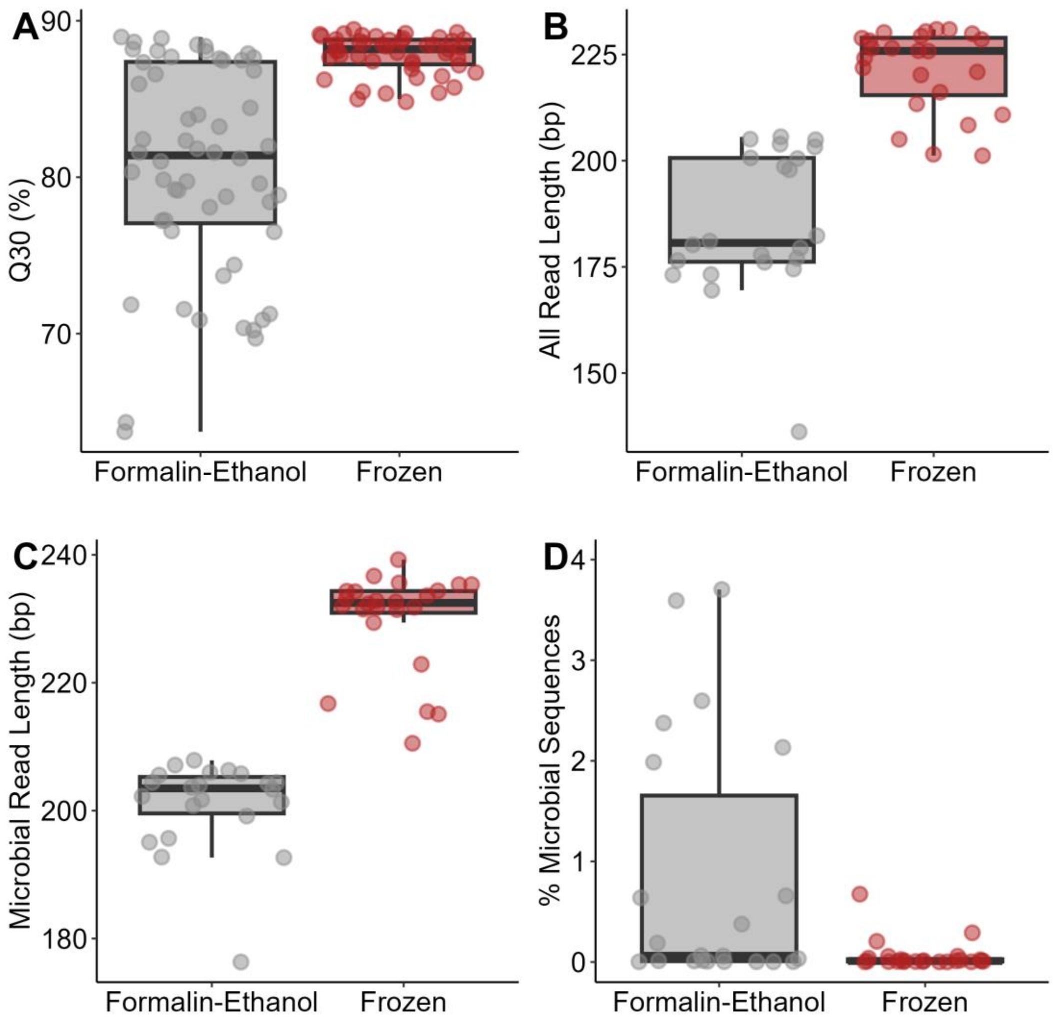


**Figure S2.** A) Quality scores (Q30); B) Merged read lengths; C) Merged microbial read length; and D) percentage microbial sequences in formalin-fixed (gray) and frozen (red) mesopelagic fish gut microbiome samples.

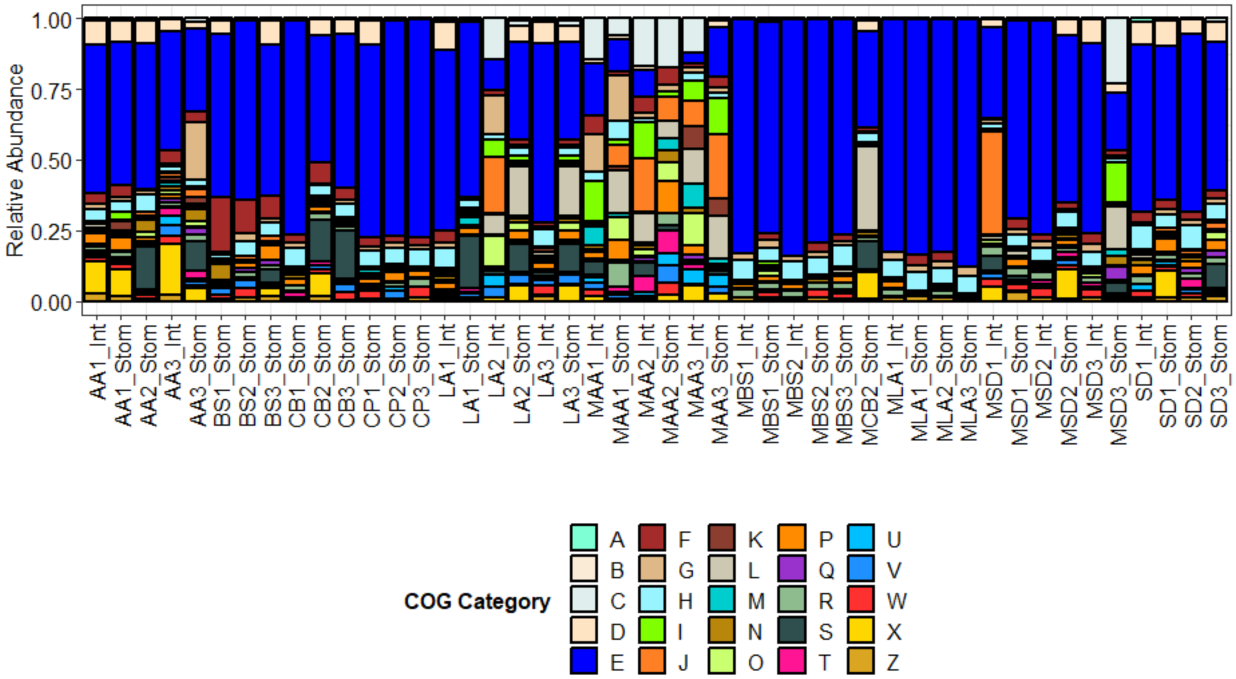


**Figure S3**. COG categories observed in mesopelagic fish gut microbiomes. [A] RNA Processing and Modification; [B] Chromatin Structure and Dynamics; [C] Energy Production and Conversion; [D] Cell Cycle Control, Cell Division, Chromosome Partitioning; [E] Amino Acid Transport and Metabolism; [F] Nucleotide Transport and Metabolism; [G] Carbohydrate Transport and Metabolism; [H] Coenzyme Transport and Metabolism; [I] Lipid Transport and Metabolism; [J] Translation, Ribosomal Structure and Biogenesis; [K] Transcription; [L] Replication, Recombination and Repair; [M] Cell Wall/Membrane/Envelope Biogenesis; [N] Cell Motility; [O] Posttranslation Modification, Protein Turnover, Chaperones; [P] Inorganic Ion Transport and Metabolism; [Q] Secondary Metabolites Biosynthesis, Transport and Catabolism; [R] General Function Prediction Only; [S] Function Unknown; [T] Signal Transduction Mechanisms; [U] Intracellular Trafficking, Secretion, and Vesicular Transport; [V] Defense Mechanisms; [W] Extracellular Structures; [X] Mobilome: Prophages, Transposons; [Y] Nuclear Structure; [Z] Cytoskeleton.

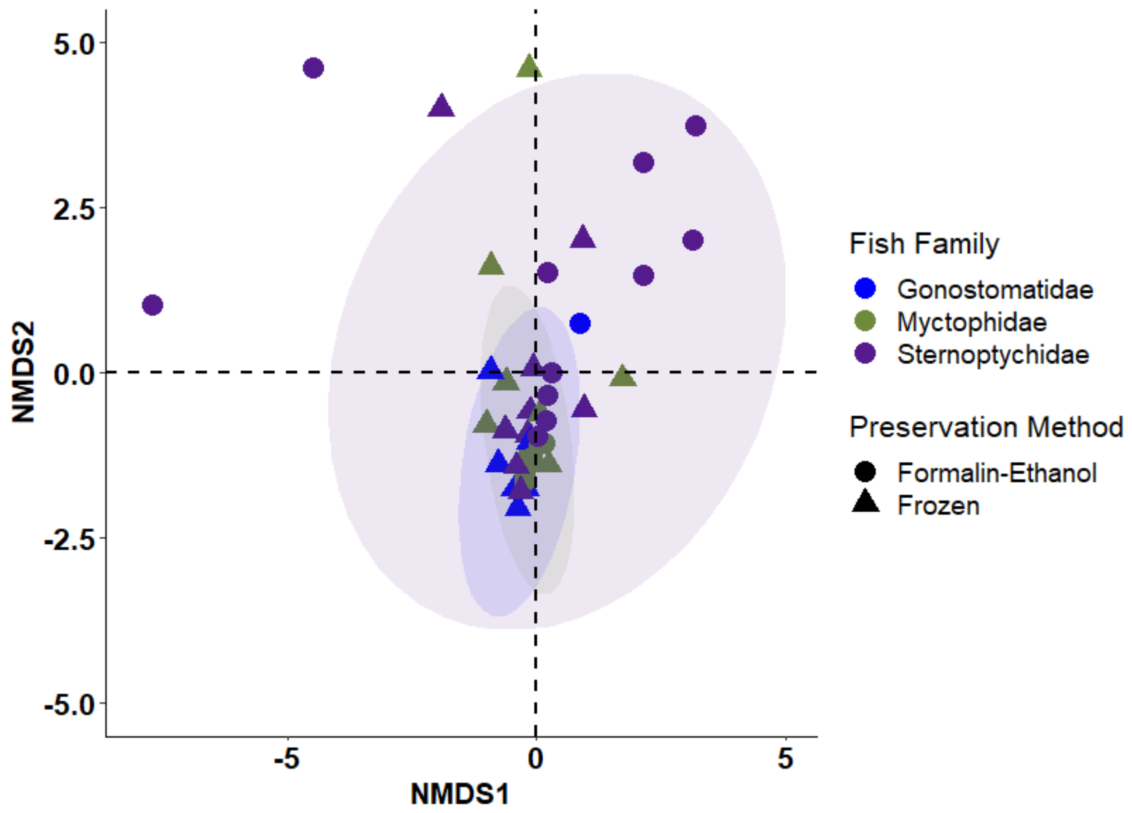


**Figure S4.** Ordination of the KOfam matrix (Bray-Curtis dissimilarity matrix of gene catalog) using non-metric multi-dimensional scaling for fish families (blue, green, purple) and preservation method (formalin-ethanol, frozen).

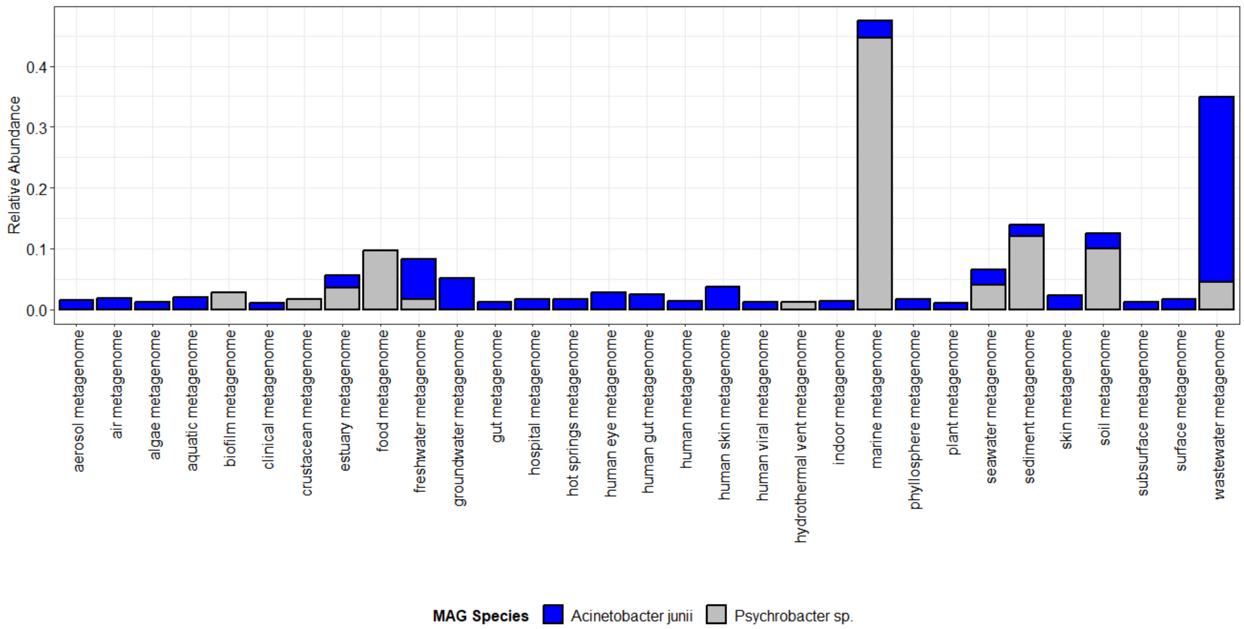


**Figure S5.** Environmental distribution of metagenome-assembled genomes (MAGs) from mesopelagic fish gut microbiomes. Blue and gray bars represent *Acinetobacter junii* and *Psychrobacter* sp., respectively.

#### Supplemental Table Captions

**Table S1.** Sample metadata for formalin-ethanol and frozen fish gut microbiomes for flow cytometric and metagenomic analyses.

**Table S2.** Pre- and postprocessing overview for formalin-ethanol and frozen fish gut microbiome samples.

### Additional Datasets

**Dataset S1.** Metagenomic gene catalog for mesopelagic fish gut microbiomes from CAZyme, COG, KOfam, and Pfam databases.

**Dataset S2.** KEGG pathway module completion for the coassembly, metagenomes, and metagenome-assembled genomes.

**Dataset S3.** Metagenome-assembled genome assembly statistics and aligned open reading frames with functional annotations from CAZyme, COG, KOfam, and Pfam databases.
